# Micronucleus Is Not a Potent Inducer of cGAS-STING Pathway

**DOI:** 10.1101/2023.10.16.561322

**Authors:** Yuki Sato, Makoto T. Hayashi

## Abstract

Micronucleus (MN) has been associated with the innate immune response. The abrupt rupture of MN membranes results in the accumulation of cGAS, potentially activating STING and downstream interferon-responsive genes. However, direct evidence connecting MN and cGAS activation has been lacking. We have developed the FuVis2 reporter system, which enables the visualization of cell nucleus carrying a single sister chromatid fusion and, consequently, MN. Using this FuVis2 reporter equipped with cGAS and STING reporters, we rigorously assessed the potency of cGAS activation by MN in individual living cells. Our findings reveal that cGAS localization to membrane-ruptured MN during interphase is infrequent, with cGAS primarily capturing MN during mitosis and remaining bound to cytosolic chromatin. We found that cGAS accumulation during mitosis neither activates STING in the subsequent interphase nor triggers the interferon response. Gamma-ray irradiation activates STING independently of MN formation and cGAS localization to MN. These results suggest that cGAS accumulation in the cytosol is not a robust indicator of its activation and that MN is not the primary trigger of the cGAS/STING pathway.

## INTRODUCTION

Micronucleus (MN), a small chromatin-containing compartment in the cytosol, is isolated from the primary nucleus (PN) and is frequently observed in aging, tumor cells, and cells exposed to genotoxic insults. Consequently, it serves as a reliable biomarker for chromosome instability (Krupina et al, 2021). MNs can form as a result of chromosome missegregation due to lagging chromosomes, acentric chromosome fragments (Fenech et al, 2011; Thompson & Compton, 2011) and breakage of anaphase chromatin bridges (Kagaya et al, 2020; Umbreit et al, 2020). Genetic material in MNs undergoes dysregulated DNA replication and DNA damage repair (Crasta et al, 2012), potentially leading to chromothripsis events (Zhang et al, 2015; Ly et al, 2016, 2019; Kneissig et al, 2019; Umbreit et al, 2020). Recently, MNs have been associated with the activation of the innate immune response through the cyclic GMP-AMP synthase (cGAS) and stimulator of interferon genes (STING) pathway (Dou et al, 2017; Glück et al, 2017; Harding et al, 2017; Mackenzie et al, 2017).

cGAS is activated by cytosolic double-stranded DNA, resulting in the production of the second messenger 2’3’-cyclic GMP-AMP (cGAMP). cGAMP is detected by STING, leading to its activation through translocation from the endoplasmic reticulum (ER) to the ER-Golgi intermediate compartment (ERGIC) and the Golgi apparatus (Hopfner & Hornung, 2020). STING subsequently activates TANK-binding kinase 1 (TBK1), which phosphorylates TBK1 itself, STING, and interferon regulatory factor 3 (IRF3) transcription factor. This cascade promotes the translocation of IRF3 into the nucleus, ultimately resulting in the activation of type I interferons and interferon-stimulated genes (Hopfner & Hornung, 2020). STING also exhibits interferon-independent activity through TBK1-dependent IκB kinase ε (IKKε) recruitment and downstream NF-κB response (Balka et al, 2020), as well as cGAS-independent non-canonical activity upon DNA damage that does not involve translocation to the Golgi (Dunphy et al, 2018). While cGAS was initially reported to reside in the cytosol to prevent self-DNA activation (Wu et al, 2013), recent studies revealed that cGAS is present not only in the cytosol (Barnett et al, 2019) but also in the nucleus during interphase (Yang et al, 2017; Gentili et al, 2019), and accumulates on mitotic chromosomes (Harding et al, 2017; Yang et al, 2017; Gentili et al, 2019; Zierhut et al, 2019). Cryo-EM structures of the cGAS-nucleosome complex have demonstrated that interaction between cGAS and histone H2A-H2B dimers sequesters the DNA-binding site of cGAS required for activation (Boyer et al, 2020; Cao et al, 2020; Kujirai et al, 2020; Michalski et al, 2020; Zhao et al, 2020). Additionally, during mitosis, hyperphosphorylation of the N-terminus disordered region of cGAS has been shown to inhibit its activation (Li et al, 2021).

It has been proposed that the nuclear membrane of MN ruptures during interphase, enabling the activation of cGAS by MN (Dou et al, 2017; Glück et al, 2017; Harding et al, 2017; Mackenzie et al, 2017; Yang et al, 2017). However, these studies relied on different cell populations to analyze cGAS localization to MN and cGAS/STING-dependent interferon responses, lacking direct evidence that MN activates cGAS-STING in the same cell. This raises questions about how cGAS can be efficiently activated by MN in the presence of suppressive chromatin-cGAS interaction, with some studies suggesting that MN may not activate cGAS (Flynn et al, 2021). Notably, irradiation, commonly used to induce MN, has been shown to cause mitochondrial DNA (mtDNA) damage and a mitochondria-dependent innate immune response (Tigano et al, 2021). These findings raise the possibility that severe genotoxic insults leading to both MN formation and mitochondrial damage may trigger mtDNA-dependent cGAS activation (Kim et al, 2023). To address whether MN is a potent activator of cGAS, a reporter system capable of inducing MN without affecting mitochondrial integrity and enabling the tracking of MN formation, cGAS localization, and STING activation in live cells is required.

We have previously developed a cell-based reporter system known as FuVis (Fusion Visualization system), which allows for the visualization of cells with defined single sister-chromatid fusions (SCF) (Kagaya et al, 2020). Live-cell imaging has demonstrated that the most prominent phenotype resulting from SCF is MN formation in subsequent cell cycles (Kagaya et al, 2020). Given that the MN induced in the FuVis system originate solely from anaphase chromatin bridges caused by SCF, the FuVis reporter provides a unique opportunity to study cGAS/STING activity upon MN formation without affecting mitochondrial function.

## RESULTS

### Second Generation of Fusion Visualization System

The first generation of the FuVis reporter system (FuVis1) comprised two distinct cell lines: FuVis-XpSIS and FuVis-XpCTRL. Both cell lines contained integrated artificial cassette sequences near telomeres on the short arm of the X chromosome, incorporating two exons (154 bp and 563 bp) of the mCitrine gene in different configurations, allowing for the detection of SCF (XpSIS) or DNA damage repair without SCF (XpCTRL) through mCitrine expression (Kagaya et al, 2020). Notably, these cell lines exhibited slight variations in morphology and growth rates, indicating potential genetic or epigenetic differences arising during the cloning process, which presented challenges in interpreting the precise effects of SCF (Kagaya et al, 2020). In response to this limitation, we aimed to develop an improved FuVis system capable of detecting both SCF and DNA damage repair distinctively in a single reporter cell line (Fig. 1A). Taking advantage of the shared N-terminus amino acid sequences between mCitrine and mCerulean3, we inserted a corresponding 3’-exon of mCerulean3 downstream of the neomycin-resistance gene (neoR) and polyA sequences within the original sister cassette sequence (Fig. 1A). By targeting spacer sequences flanking the neoR with RNA-guided endonucleases, we enabled neoR deletion, followed by mCerulean3 expression (Fig 1A, neoR deletion), as well as sporadic sister chromatid fusion, followed by mCitrine expression (Fig 1A, Sporadic sister chromatid fusion). We successfully isolated a FuVis2-XpSC33 clone that harbors a single reporter cassette integration without apparent karyotypic or growth defects (Fig. S1A-E, please refer to the Supplementary information for details).

**Figure 1.**
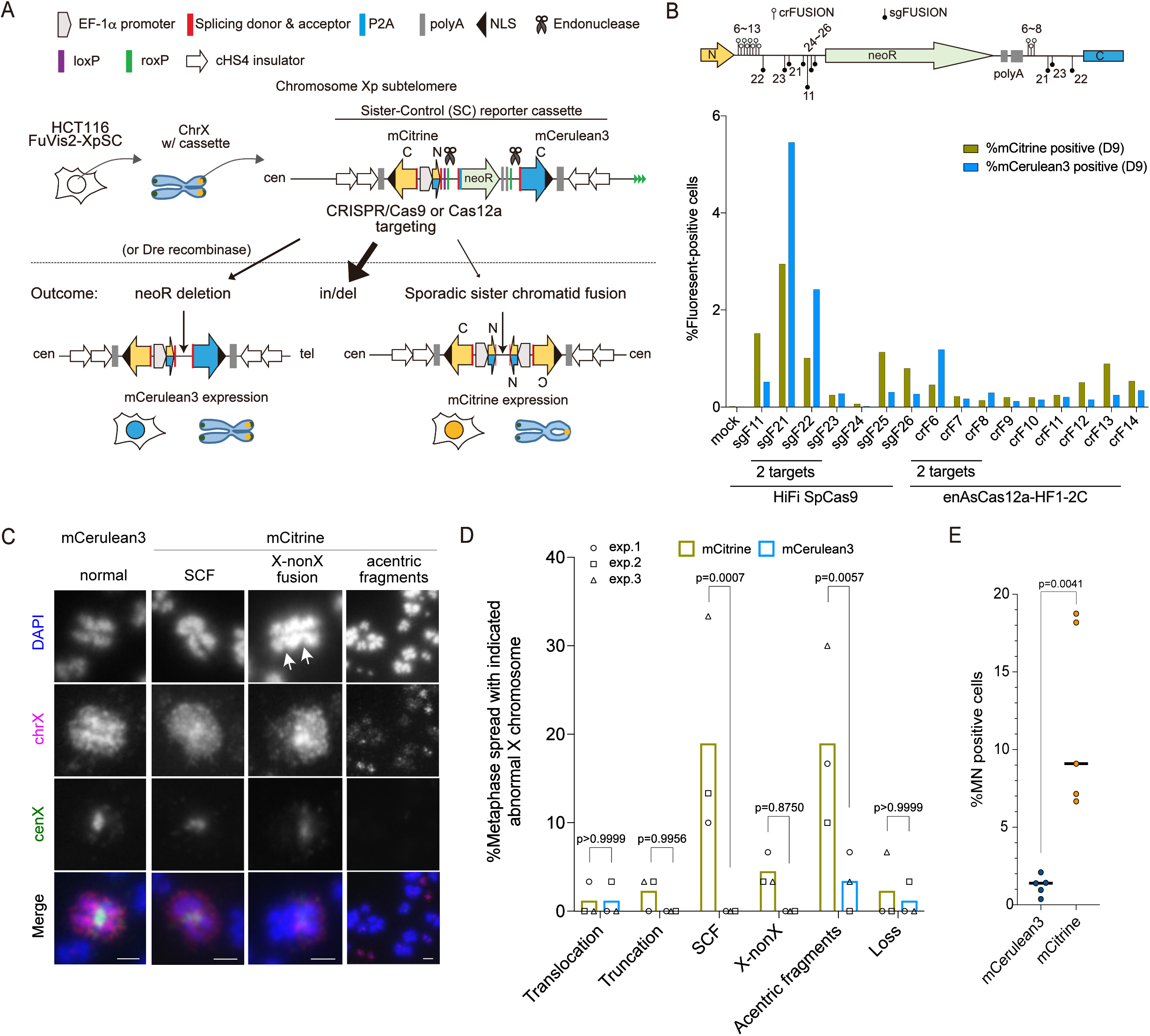
Validation of FuVis2-XpSC reporter system. (A) Schematic of the FuVis2 reporter system. The Sister-Control (SC) reporter cassette sequence, with indicated sequence features, was integrated into the subtelomere on the short arm of the X chromosome in HCT116 cells. CRISPR/Endonuclease-targeting of spacer regions flanking the neomycin-resistance gene (neoR) can result in the deletion of neoR, leading to mCerulean3 expression, or sporadic sister chromatid fusion followed by mCitrine expression. (B) Percentage of mCitrine– and mCerulean3-positive XpSC33 cells expressing the indicated endonucleases and guide RNA. The target sequence positions of each guide RNA are indicated above the graph. Cells were transduced with viruses encoding indicated endonuclease and guide RNA and analyzed at 9 days post-infection. (C) Representative images of the X chromosome in XpSC33 Cas9-sgF21 cells. At 6 days post-infection, cells were treated with 100 ng/ml colcemid for 16 hours, and mCitrine– and mCerulean3-positive cells were sorted and analyzed by chromosome spread using whole chrX (magenta) and centromere X (green) probes. Arrows indicate potential centromere loci. Scale bar, 2 µm. (D) Quantification of X chromosome abnormalities, as shown in (C), Fig. S1F and S1G. Abnormalities include translocations (non-X chromosome fragment on chrX), truncations (loss of chrX arm), SCF (sister chromatid fusion), X-nonX (presence of one cenX and one non-cenX centromere on a single chromosome), acentric fragments (small fragments of chrX without cenX signal), and loss (no chrX/cenX signal). Results from three biological replicates are shown (n = 30/experiment, ordinary one-way ANOVA followed by Sidak’s multiple comparisons). (E) Percentage of micronuclei-positive XpSC33 Cas9-sgF21 cells. At 6 days post-infection, mCerulean3– and mCitrine-positive cells were analyzed using fluorescence microscopy. Results from five biological replicates are shown (n = at least 15/experiment for mCitrine and 216/experiment for mCelulean3, two-tailed Student’s t-test).

To validate the FuVis2 reporter, we targeted various sequences flanking neoR using two endonucleases: the SpCas9 variant HiFi SpCas9 (Cas9(HiFi)) and AsCas12a variant enAsCas12a-HF1-2C (Cas12a(HF1)) (Vakulskas et al, 2018; Kleinstiver et al, 2019). Guide RNAs (sgFUSION and crFUSION) were designed for both endonucleases to target either a single site upstream of neoR or two sites flanking neoR (Fig. 1B). XpSC33 cells were transduced with a virus encoding either Cas9(HiFi)-sgFUSION (sgF) or Cas12a(HF1)-crFUSION (crF) and analyzed on day 9 using flow cytometry. Among these constructs, only guide RNAs targeting two neoR-flanking sequences (sgF21, sgF22, and crF6) induced both mCerulean3 and mCitrine expression (Fig. 1B). Guide RNAs targeting a single site (sgF11, sgF25, sgF26, crF12, crF13, and crF14) induced mCitrine expression with a background level of mCerulean3 expression (Fig. 1B). For subsequent analysis, we selected Cas9(HiFi)-sgF21 (hereafter Cas9-sgF21), which induced the highest levels of both mCitrine and mCerulean3.

To analyze X chromosome abnormalities, mCitrine– and mCerulean3-positive XpSC33 Cas9-sgF21 cells were sorted and subjected to dual-colored FISH analysis using whole X chromosome painting (ChrX) and chromosome X centromere specific (cenX) probes.

Compared to mCerulean3-positive cells, mCitrine-positive cells exhibited a significantly increased rate of abnormal X chromosomes, including SCF and acentric fragments (Fig. 1C, D, and S1F, G). While a slight increase in chromosome fusion between X and non-X chromosomes was also observed, it did not reach statistical significance (Fig. 1C, D). Time-course analysis of XpSC33 cells expressing different endonuclease and guide RNA pairs showed that mCerulean3-positive cells reached a plateau as early as 6 days post-infection, while mCitrine-positive cells peaked around day 6 and gradually decreased, irrespective of the efficiency of the endonucleases and guide RNA used (Fig. S1H). This kinetic pattern aligns with the assumption that a single mCitrine gene locus generated by SCF can be transmitted to either one of two daughter cells, resulting in the gradual loss of mCitrine protein in the other lineage that did not inherit the mCitrine gene (Kagaya et al, 2020). Importantly, mCitrine-positive cells exhibited increased MN formation compared to mCerulean3-positive cells 6 days post-infection (Fig. 1E). These findings are consistent with previous results obtained from the FuVis1 system, confirming that a single SCF event can lead to MN formation.

### Sister chromatid fusion causes micronuclei following the first mitosis

To investigate the kinetics of MN formation in the FuVis2 system, we conducted live-cell imaging using XpSC33 Cas9-sgF21 cells. During the first interphase when cells became fluorescent-positive, neither mCitrine-nor mCerulean3-positive cells displayed MN (Fig 2A-C). However, in the second interphase, 40% of mCitrine-positive cells developed MN, while mCerulean3-positive cells remained MN-free (Fig. 2C). This result further supports the notion that MN originates from a single SCF event that experienced breakage during the first mitotic exit.

**Figure 2.**
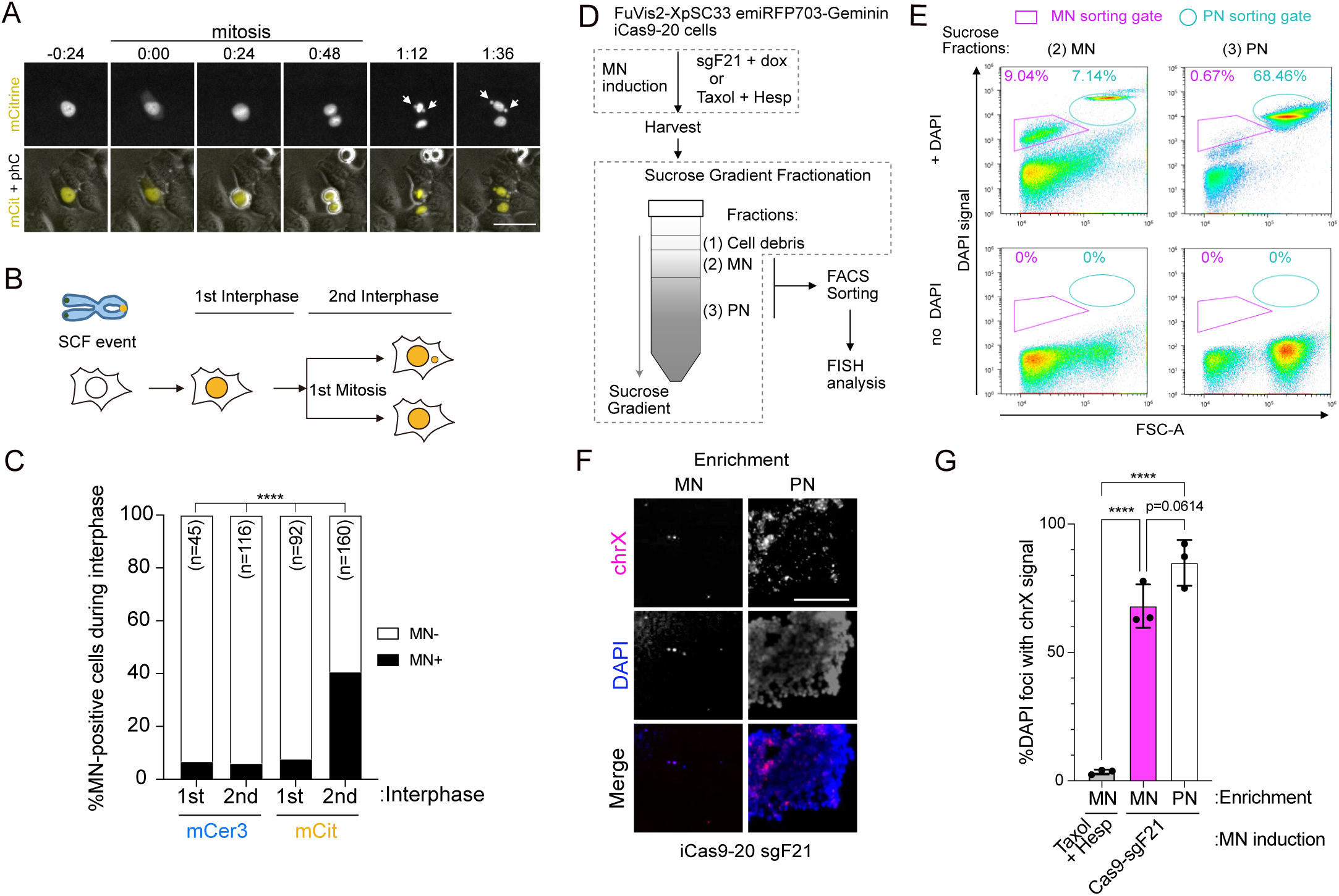
Sister chromatid fusion causes micronuclei following the first mitosis. (A) Representative live-cell images of micronuclei (MN) formation in mCitrine-positive XpSC33 Cas9-sgF21 cells. White arrows indicate MN. Scale bar, 25 µm. phC, phase contrast. (B) Schematic of the cell cycle counts following SCF event. (C) Percentage of MN-positive cells at the indicated cell cycle stages shown in (B). mCerulean3 (mCer3)– and mCitrine (mCit)-positive XpSC33 Cas9-sgF21 cells were analyzed by live-cell imaging from 4 days to 7 days post-infection. Data was collected from three biological replicates (Chi-square test). ****p<0.00001. (D) Schematic showing the enrichment of MN and primary nuclei (PN). XpSC33 emiRFP703-Geminin iCas9-20 cells were transduced with sgF21-encoding virus and exposed to 0.1 µg/ml dox for 8 days or treated with 250 nM Taxol and 250 nM Hesperadin for 2 days. Cell extracts were fractionated by sucrose gradient to obtain MN and PN fractions. Each fraction was DAPI-stained and subjected to FACS sorting to further enrich MN and PN, as shown in (E). Enrichment of MN and PN was confirmed by FISH analysis, as shown in (F). (E) Representative FACS analysis of MN and PN fractions from the sucrose gradient with or without DAPI staining. (F) Representative FISH images of MN– and PN-enriched samples from XpSC33 emiRFP703-Geminin iCas9-20 sgF21 cells. The sorted samples from (E) were further enriched by centrifugation and analyzed using the whole chrX probe (magenta). Scale bar, 10 µm. (G) Percentage of DAPI foci with chrX signals, as shown in (F). Results from three biological replicates are shown (n = at least 70/experiment, ordinary one-way ANOVA followed by Tukey’s multiple comparisons). ****p<0.00001.

The continuous expression of Cas9 raises concerns about potential off-target genomic damage, which could lead to unintended MN formation. To address this concern, we isolated a clone of XpSC33 cells equipped with a doxycycline (dox)-inducible Cas9(HiFi), subsequently renamed as XpSC33-iCas9-20 (Fig. S2A-F, please refer to the Supplementary information for details). XpSC33-iCas9-20 cells were transduced by the sgF21-encoding virus in the presence of 0.1 µg/ml dox for 1 day and analyzed from day 2 to 6 using a flow cytometer, confirming the expected expression of both mCitrine and mCerulean3 (Fig. S3A). Live-cell analysis revealed a significant increase in MN-positive cells during the second interphase among mCitrine-positive, but not mCerulean3-positive, cells (Fig. S3B).

To further validate the nature of MN, we aimed to purify SCF-derived MN from XpSC33 cells. Since MN isolation requires a sufficient number of cells, and mCitrine-positive cells are rare, we decided to use the entire population of sgF21 expressing XpSC33 iCas9-20 cells. However, live-cell analysis revealed that both mCitrine– and mCerulean3-positive populations exhibited MN-positive cells in the first interphase (Fig. 2C and S3B), likely stemming from background MN formation unrelated to the SCF event. Since these cells did not divide frequently, we attempted to collect a cycling population to accumulate cells with SCF-derived MN. For this purpose, XpSC33 iCas9-20 cells were transduced with a virus encoding emiRFP703-Geminin, a derivative of the FUCCI reporter system for visualizing the S/G2/M phase of the cell cycle (Sakaue-Sawano et al, 2008). Transduced cells were sequentially sorted twice to enrich cells with the expected reporter expression, validated by aphidicolin treatment and serum starvation (Fig. S3C). The resulting XpSC33 emiRFP703-Geminin iCas9-20 cells were transduced with the sgF21-encoding virus, and emiRFP703-positive cells were sorted on day 8 post-infection. Cell extracts were subjected to sucrose-gradient fractionation and sorting by DAPI-staining for MN and Primary Nuclei (PN) purification (Fig. 2D, E). The resulting MN– and PN-enriched samples were subjected to FISH analysis using the ChrX probe. As anticipated, the PN-enriched sample consistently exhibited ChrX focus formation (Fig. 2F and 2G). Remarkably, we found that the MN-enriched sample was very frequently painted with the ChrX probe (Fig. 2F and 2G). In contrast, a similar painting was not observed in a MN-enriched sample from cells treated with a microtubule stabilizer Taxol and Aurora kinase B inhibitor Hesperadin for 48 hours (Fig. 2D, G). Collectively, these results suggest that the SCF-derived chromatin bridge of X chromosomes is disrupted during the first mitosis, leading to MN formation in the subsequent cell cycle. Thus, the FuVis2 reporter system offers a unique opportunity to explore the fate of MN originating solely from a single SCF event on the short arm of the X chromosome.

### MN-derived chromatin is captured by cGAS upon mitotic nuclear envelope breakdown

Previous studies have suggested that the MN membrane ruptures during interphase, leading to the accumulation and activation of cGAS (Dou et al, 2017; Harding et al, 2017; Mackenzie et al, 2017). We refer to this phenomenon as ‘interphase-cGAS accumulation in MN’ or ‘i-CAM’ and aimed to determine the frequency of i-CAM in XpSC33 Cas9-sgF21 cells expressing mScarlet-cGAS. A long-term live-cell analysis of mCitrine-positive cells revealed that i-CAM is a rare event, occurring in only 6.5% of MN-positive cells (Fig. 3A). Instead, we observed unique cGAS localization patterns during mitosis, which could be classified into three categories. First, in MN-negative cells and 9.5% of MN-positive cells, mitotic cGAS localized to PN-derived chromosomes, consistent with previous reports (Harding et al, 2017; Gentili et al, 2019; Zierhut et al, 2019)(Fig. 3B, C; PN only). Second, in 47.6% of MN-positive cells, cGAS localized to both MN– and PN-derived chromosomes (Fig. 3B, C; PN+MN). Lastly, in 42.9% of MN-positive cells, cGAS robustly accumulated in the MN-derived chromosome region upon nuclear envelope breakdown (NEBD) (Fig. 3B-D; MN only). Collectively, these findings revealed that 90.5% of MN-positive cells that entered mitosis exhibited mitotic cGAS accumulation in MN-derived chromatin, which we term ‘m-CAM’ (Fig. 3E).

**Figure 3.**
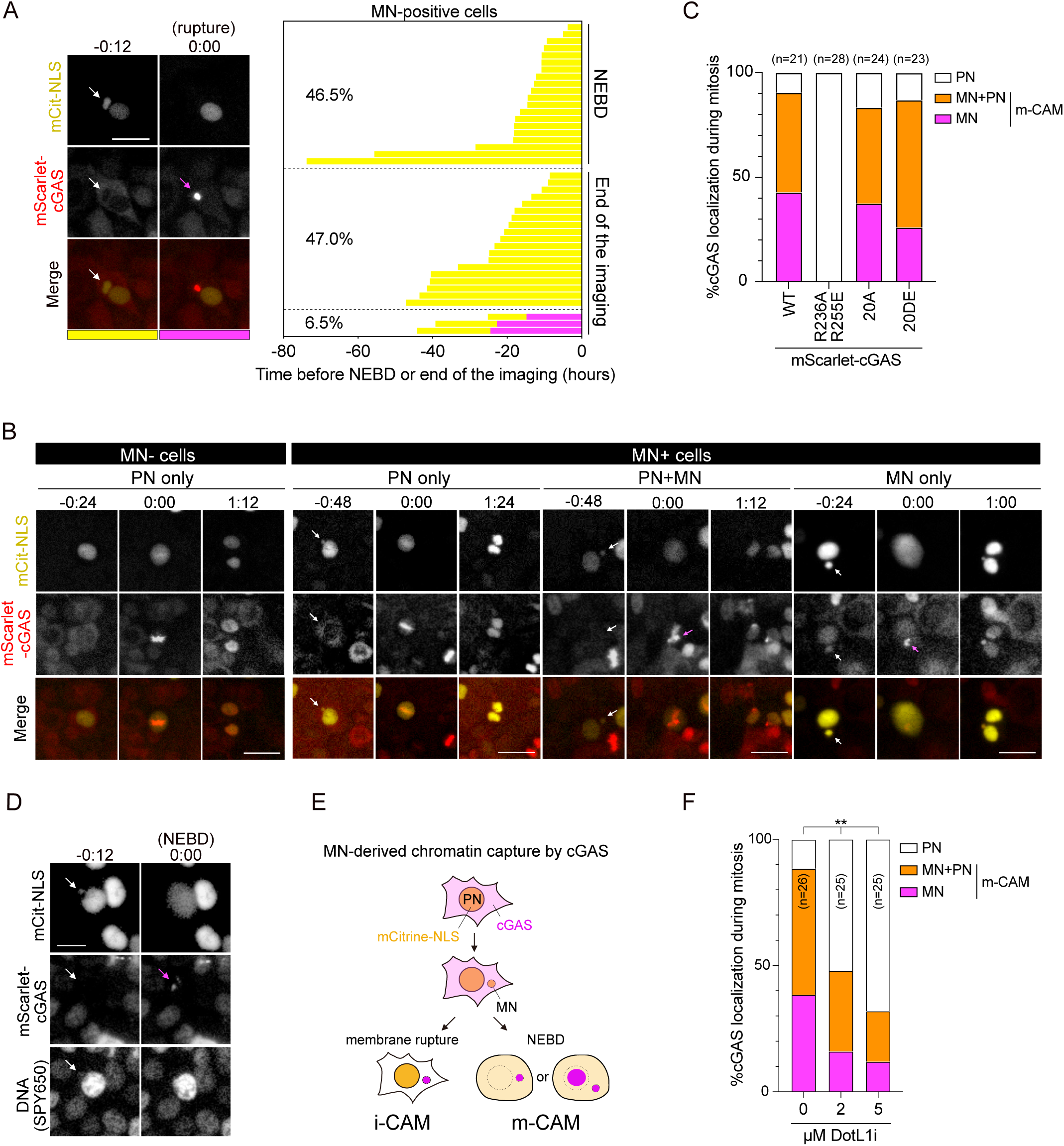
SCF-derived MN is captured by cGAS upon mitotic nuclear envelope breakdown. (A) Live-cell analysis of MN captured by cGAS during interphase in XpSC33 mScarlet-cGAS Cas9-sgF21 cells. Left: a representative MN membrane rupture event is shown. White arrows indicate MN with intact membrane, as evidenced by mCitrine-NLS localization; magenta arrow indicates cGAS accumulation in MN. Scale bar, 25 µm. Right: cGAS localization pattern in XpSC33 mScarlet-cGAS Cas9-sgF21 live-cell imaging. Each bar represents mCitrine– and MN-positive cell as it progresses through interphase to mitosis (NEBD, nuclear envelope breakdown) or the end of imaging, both of which are set as T = 0. Yellow bars, mCitrine-positive and mScarlet-negative MN; magenta bars, mCitrine-negative and mScarlet-positive MN, as shown on the left. Percentage of each category is shown. (B) Representative live-cell images of mScarlet-cGAS localization upon NEBD in MN-negative and MN-positive XpSC33 mScarlet-cGAS Cas9-sgF21 cells. NEBD was recognized by the diffusion of mCitrine-NLS signal. White arrows indicate MN with intact membrane before NEBD; magenta arrows indicate mScarlet-cGAS foci on MN-derived chromatin upon NEBD. Scale bar, 25 µm. (C) Percentage of cGAS localization patterns upon NEBD as shown in (B). MN-positive XpSC33 Cas9-sgF21 cells expressing WT or indicated mScarlet-cGAS mutants were analyzed by live-cell imaging from 4 days to 7 days post-infection of Cas9-sgF21. (D) Representative images of mScarlet-cGAS accumulation in MN-derived DNA. XpSC33 mScarlet-cGAS Cas9-sgF21 cells were incubated with SPY650 for 2 hours and analyzed by live-cell imaging. White arrows indicate MN with intact membrane; magenta arrow indicates mScarlet-cGAS accumulation in MN-derived DNA locus. Scale bar, 10 µm. (E) Schematic illustrating two distinct pathways of cGAS in the initial capture of MN: i-CAM (interphase cGAS accumulation in MN, initiated by MN membrane rupture event) and m-CAM (mitotic cGAS accumulation in MN-derived chromatin, initiated by NEBD). (F) Percentage of cGAS localization patterns upon NEBD as shown in (B). XpSC33 mScarlet-cGAS cells were treated with the indicated dose of SGC0946, a DotL1 inhibitor, for 1 week and transduced with Cas9-sgF21-encoding virus. MN-positive cells were analyzed by live-cell imaging from 4 days to 7 days post-infection of Cas9-sgF21. Data were collected from two biological replicates (Chi-square test between m-CAM and no m-CAM) **p<0.001.

We further tracked the reformation of MN and cGAS localization in the subsequent interphase, categorizing them into four groups (Fig. S3D): (1) mCitrine-positive MN with cGAS accumulation, (2) mCitrine-negative MN with cGAS accumulation, (3) mCitrine-positive MN without cGAS accumulation, and (4) no evidence of MN. Notably, we observed that cGAS accumulated in MN in approximately half of the m-CAM-derived MN-positive cells (Fig. S3D). This observation aligns with previous reports demonstrating cGAS accumulation in MN among fixed interphase cells (Dou et al, 2017). Our results suggest that the cGAS accumulation in MN observed in fixed cells mainly arises from MN that has experienced the m-CAM event.

To explore the mechanism behind m-CAM, XpSC33 Cas9-sgF21 cells expressing three cGAS mutants were subjected to live-cell analysis. We discovered that cGAS^R236A-R255E^ mutant, which carries mutations on the nucleosome-binding surface (Volkman et al, 2019), completely abolished the m-CAM event while retaining localization to PN-derived chromosomes (Fig. 3C and S3E). On the other hand, no effect on m-CAM was observed in cells expressing phosphomimetic (cGAS^20DE^) and phosphor-null (cGAS^20A^) mutants of its N-terminal domain, which harbor mutations in twenty Ser/Thr sites required for mitotic inactivation of cGAS (Li et al, 2021) (Fig. 3C and S3E). This result indicates that the nucleosome binding ability of cGAS is crucial for m-CAM, which is distinct from mitotic cGAS localization to PN-derived chromosomes. We further addressed if m-CAM is influenced by modifying MN-specific histone modification, H3K79me2, known to recruit cGAS to interphase MN (MacDonald et al, 2023). Pre-treatment with a DOT1L inhibitor SGC0946 for seven days, which abolishes H3K79me2 (MacDonald et al, 2023), significantly suppressed the m-CAM event (Fig. 3F). These results suggest that the H3K79m2 mark on MN allows cGAS to interact more efficiently with nucleosome upon mitotic entry.

### m-CAM does not lead to STING activation

The dominance of the m-CAM event and the persistence of cytoplasmic cGAS foci in the subsequent interphase raised a possibility that STING is activated in the subsequent cell cycle. Since TBK1 and IRF3 can be activated independently of cGAS-STING pathways (Liu et al, 2015), we aimed to directly monitor the activity of STING. To achieve this, XpSC33 cells were transduced with viruses encoding emiRFP703-cGAS and mRuby3-STING reporters (Balka et al, 2023; Kuchitsu et al, 2023). STING translocates from the ER to the Golgi apparatus during activation (Mukai et al, 2016). Consistently, mRuby3-STING accumulated at the Golgi apparatus 2 hours after exposure to compound 3, a potent STING agonist (Ramanjulu et al, 2018)(Fig. 4A). We utilized the maximum intensity and average intensity of mRuby3-STING in a cell to assess STING accumulation as an indicator of its activation (Fig. 4A, STING Accumulation Index: St-AI). To validate the reliability of St-AI, cells were immunostained for pSTING-S366, a TBK1-dependent phosphorylation indicative of its activation (Liu et al, 2015)(Fig. 4B). Based on the scatter plot of pSTING-S366 intensity and St-AI, we observed a strong correlation between St-AI values and pSTING-S366 signal intensities (Fig. 4C and S4A). We defined St-AI values greater than 2.0 as indicative of STING activation (Fig. 4C). Serial dilution of compound 3 showed that pSTING-S366 intensity and St-AI exhibited a similar threshold concentration for indicating STING activation (Fig. 4D, S4B, and S4C), which correlated well with the upregulation of cxcl10, an interferon gamma-induced inflammatory marker (Fig. 4E). Time-lapse analysis confirmed that, compared to the mock control, St-AI gradually increased after the transfection of pMAX-TurboGFP (GFP) plasmid as a source of cytosolic dsDNA (Fig. 4F, G, and S4D). shRNA knockdown of cGAS completely abolished the increase in St-AI following pMAX-GFP transfection but not compound 3 (Fig. 4H, S4E, and S4F), confirming cGAS-dependent STING activation in the presence of cytosolic dsDNA. The attenuation of St-AI by shcGAS under the compound 3 condition may be attributed to the loss of secondary activation of the cGAS-STING cycle caused by dsDNA released from dead cells (Messaoud-Nacer et al, 2022). In conclusion, we consider St-AI a valuable indicator of STING activation in live cells.

**Figure 4.**
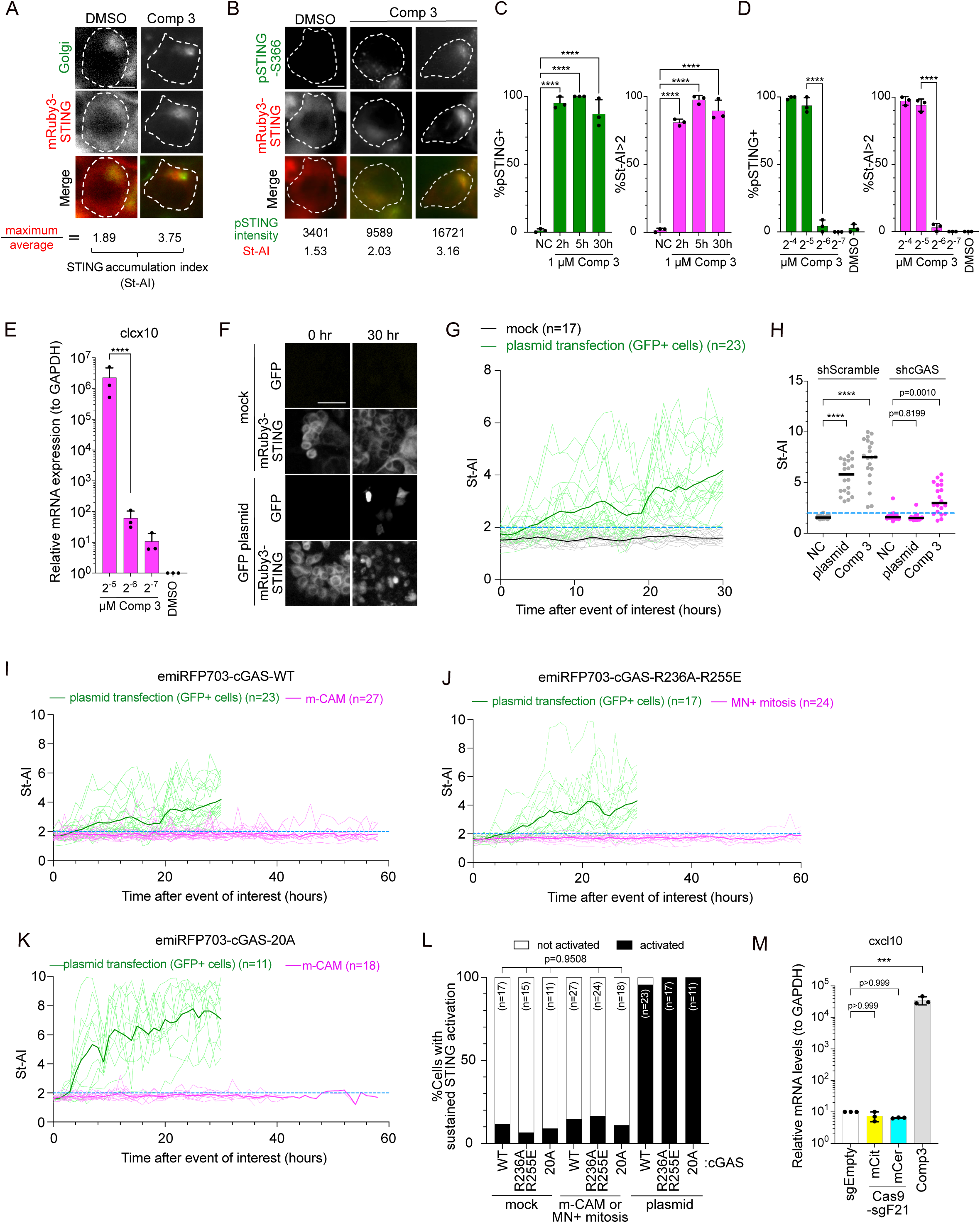
m-CAM does not lead to STING activation in subsequent interphase. (A) Representative images of mRuby3-STING and Golgi apparatus colocalization upon STING activation. XpSC33 emiRFP703-cGAS mRuby3-STING cells were treated with 1 µM compound 3 for 2 hours and stained with the Cell Navigator NBD Ceremide Golgi Staining Kit (green). Dot lines indicate the cell membrane determined by phase-contrast images. Maximum and average intensities of mRuby3-STING inside a cell membrane were used to calculate STING accumulation index (St-AI) as shown below. Scale bar, 10 µm. (B) Representative images of mRuby3-STING and pSTING-S366 colocalization upon its activation. XpSC33 emiRFP703-cGAS mRuby3-STING cells were treated with 1 µM compound 3 for 2 hours, fixed and stained with anti-pSTING-S366 (green). Dot lines indicate the cell membrane. Total pSTING-S366 intensity (arbitrary units) inside a cell membrane and St-AI are shown below. Scale bar, 10 µm. (C) Percentage of pSTING-S366 positive cells (left) and cells with St-AI greater than 2.0 (right) in individual XpSC33 emiRFP703-cGAS mRuby3-STING cells treated with 1 µM compound 3 for the indicated hours. Cells were fixed and stained for pSTING-S366 as shown in (B). pSTING-S366 intensities over 6,000 (arbitrary units) were defined as pSTING-S366 positive (Fig. S4A). Results from three biological replicates are shown (n = at least 35/experiment, ordinary one-way ANOVA followed by Tukey’s multiple comparisons test). ****p<0.00001. (D) Percentage of pSTING-S366 positive cells (left) and cells with St-AI greater than 2.0 (right) at indicated doses of compound 3. XpSC33 emiRFP703-cGAS mRuby3-STING cells were treated with compound 3 for 5 hours and analyzed as in (B). Results from three biological replicates are shown (n = at least 38/experiment, ordinary one-way ANOVA followed by Tukey’s multiple comparisons test). ****p<0.00001. (E) Relative mRNA levels of cxcl10 normalized to GAPDH. XpSC33 emiRFP703-cGAS mRuby3-STING cells were treated with indicated doses of compound 3 for 5 hours and harvested 25 hours after washout of compound 3. Total RNA was subjected to RT-qPCR (n = 3 biological replicates, unpaired t-test). ****p<0.00001. (F) Representative images of GFP and mRuby3-STING at 30 hours post-transfection of the pMAX-TurboGFP plasmid. XpSC33 emiRFP703-cGAS mRuby3-STING cells were transfected with pMAX-TurboGFP and subjected to live-cell imaging. Scale bar, 50 µm. (G) Time-course analysis of St-AI after plasmid transfection as shown in (F). Bold lines indicate the average of individual cells in each condition. Only GFP-positive cells were analyzed after transfection. The blue dot line represents St-AI = 2.0. (H) St-AI in XpSC33 emiRFP703-cGAS mRuby3-STING cells expressing shScramble or shcGAS (n = 20, ordinary one-way ANOVA followed by Tukey’s multiple comparisons test). shRNA-transduced cells were either transfected with pMAX-TurboGFP for 30 hours or exposed to 1 µM compound 3 for 30 hours. The blue dot line represents St-AI = 2.0. (I-K) Time-course analysis of St-AI after pMAX-TurboGFP plasmid transfection or the m-CAM event in XpSC33 mRuby3-STING cells expressing indicated cGAS variants with miRFP670-tag. Only GFP-positive cells were analyzed after transfection. The blue dot line represents St-AI = 2.0. (J) Since emiRFP703-cGAS^R236A-R255E^ does not show m-CAM, cells entering mitosis with MN were analyzed in the following interphase. (L) Percentage of sustained STING activation (St-AI > 2.0 for a duration over 4 hours) in cells expressing indicated cGAS mutants from (I-K). (M) Relative mRNA levels of cxcl10 normalized to GAPDH. mCerulean3– and mCitrine-positive XpSC33 Cas9-sgF21 cells were sorted at 7 days post-infection. XpSC33 Cas9-sgEmpty cells and XpSC33 cells treated with 1 µM compound 3 for 5 hours followed by 25 hours recovery were harvested as negative and positive control, respectively. Total RNA was subjected to RT-qPCR (n = 3 biological replicates, ordinary one-way ANOVA followed by Tukey’s multiple comparisons test). ***p<0.0001.

To address STING activation after m-CAM, we performed live-cell imaging in XpSC33 emiRFP703-cGAS mRuby3-STING cells transduced with the Cas9-sgF21-encoding virus. We first confirmed that lentivirus transduction itself does not activate STING (Fig. S4G), and that the m-CAM event is dominant over the i-CAM event under these conditions as well (Fig. S4H). Time-course analysis revealed that St-AI remained unchanged during the interphase following the m-CAM event (Fig. 4I). Since both nucleosome binding and mitotic hyper-phosphorylation attenuate cGAS activation (Volkman et al, 2019; Li et al, 2021), we performed the same experiments in cells expressing emiRFP703-cGAS^R236A-R255E^ and emiRFP703-cGAS^20A^ (Fig. 4J and 4K). To compare the STING activation rates under various live-cell imaging conditions, we defined sustained STING activation as St-AI values exceeding 2.0 for a duration over 4 hours during a specified interphase. We found no significant increase in sustained STING activation after m-CAM, compared to the plasmid transfection control, in cells expressing not only cGAS-WT but also –R236A-R255E and – 20A (Fig. 4L). We attempted but failed to obtain XpSC33 cells expressing emiRFP703-cGAS^R236A-R255E-20A^ mutant due to strong toxicity (Li et al, 2021). These results suggest that cGAS activation is strongly suppressed during and after the m-CAM event. In agreement with this result, neither mCitrine-positive nor mCerulean3-positive SC33 Cas9-sgF21 cells showed any induction of cxcl10 (Fig. 4M), suggesting that even faint activation of cGAS was absent from the mCitrine-positive population.

### STING activation after irradiation is independent of MN formation

To clarify the reasons for discrepancies between our findings and prior reports (Dou et al, 2017; Glück et al, 2017; Harding et al, 2017; Mackenzie et al, 2017), we assessed St-AI following MN formation induced by gamma-ray irradiation. XpSC33 emiRFP703-cGAS mRuby3-STING cells were transduced with a virus encoding full-length mCitrine-NLS to visualize nuclei, irradiated at 1 Gy, and subjected to live-cell imaging. As expected, irradiated cells exhibited MN as cytosolic mCitrine foci following the first mitosis post-irradiation (Fig. 5A, B, and S5A), which is comparable to SCF-induced MN formation (Fig. 2C). Initially, we examined the cGAS localization pattern to MN and observed that only 10.3% and 9.4% of MN-positive cells exhibited the i-CAM event during the 2^nd^ and 3^rd^ interphase, respectively, while 77.8% and 92.3% of cells that entered mitosis displayed the m-CAM event in the 2^nd^ and 3^rd^ mitosis, respectively (Fig. 5C). These results suggest that m-CAM is common in the initial MN capture by cGAS. Subsequently, we analyzed St-AI during interphase following i-CAM and m-CAM events. Among 17 i-CAM events observed, 11 cells did not show St-AI increase after the i-CAM event (Fig. S5B, C). Two cells showed a sharp St-AI increase after the i-CAM event (Fig. S5D), and four did not show such a spike but sustained STING activation (Fig. S5E, F). However, among the six cells that exhibited STING activation, five of them showed sustained STING activation before the i-CAM event (Fig. S5D, E). This result suggests that i-CAM has a potential to trigger STING activation, but in most cases, it is not sufficient, and STING is activated by other stimuli. In agreement with this assumption, both the MN-negative lineage and the interphase following m-CAM exhibited a similar increase in St-AI (Fig. 5A, D-F, and S5G), suggesting that STING is activated irrespective of MN formation after 1 Gy IR exposure. Cells expressing emiRFP703-cGAS^R236A-R255E^ exhibited an increased frequency of sustained STING activation in both MN-negative and MN-positive lineages (Fig. 5G-I), suggesting that nucleosomal DNA leaked into cytoplasm, which could not be visualized by mCitrine-NLS nor emiRFP703-cGAS, inhibited cGAS activation in irradiated cells.

**Figure 5.**
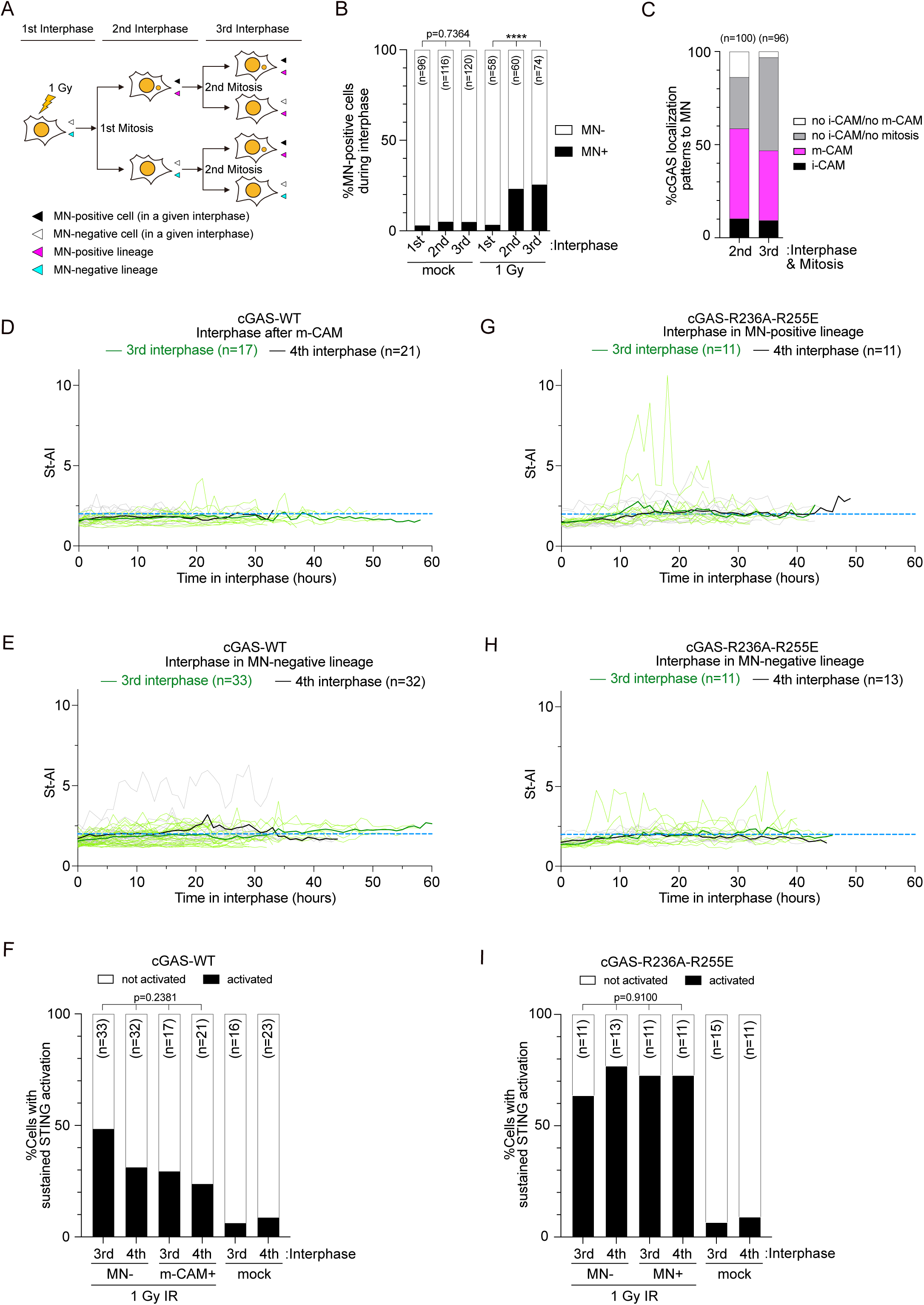
MN independence in STING activation after irradiation. (A) Schematic of cell cycle tracking after irradiation. In this live-cell imaging analysis, cells with and without mCitrine-positive MN are identified as MN-positive and MN-negative cells, respectively, in each interphase. All descendant cells originating from a MN-positive cell are defined as the MN-positive lineage. Note that a cell in the MN-positive lineage can become MN-negative in different interphases, while all cells in the MN-negative lineage remain MN-negative throughout. (B) Percentage of MN-positive cells at different cell cycle stages post-irradiation. Data were collected from four biological replicates (Chi-square test). XpSC33 mCitrine-NLS emiRFP703-cGAS mRuby3-STING cells were exposed to 1 Gy IR and subjected to live-cell imaging. ****p<0.00001. (C) Percentage of cGAS localization patterns in MN at the indicated cell cycle stages post-irradiation. MN-positive cells were tracked for emiRFP703-cGAS accumulation. No i-CAM/no m-CAM indicates MN-positive cells entered mitosis without exhibiting any signs of m-CAM. (D-E) Time-course analysis of St-AI following the m-CAM event (D) or in MN-negative lineage (E) at the indicated cell cycle stages post-irradiation, as shown in (A). (F) Percentage of sustained STING activation (St-AI > 2.0 for a duration over 4 hours) in D and E. (G-H) Time-course analysis of St-AI in indicated count of the cell cycle post-irradiation in XpSC33 mCitrine-NLS emiRFP703-cGAS^R236A-R255E^ mRuby3-STING cells. Results from cells entering mitosis with MN (G) and MN-negative lineages (H) are shown. Percentage of sustained STING activation (St-AI > 2.0 for a duration over 4 hours) in G and H.

## DISCUSSION

In this study, we aimed to rigorously evaluate the potency of MN as an activator of the cGAS-STING pathway. Our FuVis2 reporter system allows the visualization of the nucleus in cells that have acquired a single SCF on the X chromosome, serving as an ideal reporter to assess cGAS-STING activity after MN formation without compromising mitochondrial integrity. Importantly, MN is almost exclusively derived from chromosome fusion in this reporter, which emulates MN formation in the early tumorigenesis stage called telomere crisis (Nassour et al, 2019).

We have successfully introduced cGAS and STING reporters into the FuVis2 reporter cells and confirmed that the accumulation of STING quantified as the St-AI provides a good-quality indicator of STING activation, which is validated by pSTING-S366 and downstream cxcl10 expression. Our live-cell data suggest that chromosomes in MN can be captured by cGAS in interphase and mitosis through nuclear envelope rupturing and NEBD, respectively. In contrast to previous reports that emphasized the former i-CAM event (Harding et al, 2017; Mackenzie et al, 2017), our results suggest that the primary pathway of MN-chromatin detection by cGAS is through the latter m-CAM event, which depends on the nucleosome-binding motif of cGAS and histone H3K79me2-mediated exposure of the cGAS-interacting acidic patch of H2A-H2B. This mechanism is distinct from cGAS localization to PN-derived chromosomes during mitosis, which may depend on DNA-binding surfaces residing in K173-I220 and H390-C405 in cGAS (Gentili et al, 2019).

Although about one third of post-m-CAM G1 cells exhibited cytoplasmic cGAS foci formation, St-AI analysis indicated that m-CAM does not lead to activation of cGAS and STING in the following interphase. Neither STING activation nor cxcl10 expression was observed in mCitrine-positive XpSC33 Cas9-sgF21 cells, suggesting that, contrary to the previous report (Flynn et al, 2021), not only MN but also chromatin bridges caused by SCF do not activate cGAS efficiently. The complete absence of cxcl10 upregulation, a highly sensitive marker of the interferon response, in the mCitrine-positive XpSC33 Cas9-sgF21 cells suggests that there is not even a faint activation of cGAS in this population. Moreover, neither cGAS^R236A-R255E^ nor cGAS^20A^ mutants could activate STING after m-CAM. It is less likely that the emiRFP703-tag abolished cGAS^R236A-R255E^ enzymatic activity, since emiRFP703-cGAS^R236A-R255E^ expressing cells showed increased STING activation after irradiation. We assume that both nucleosome-binding and N-terminus hyper-phosphorylation mechanisms, as well as other inhibitory mechanisms including BAF and TREX1 (Guey et al, 2020; Mohr et al, 2021), redundantly suppress cGAS activation upon MN formation. Although these possibilities need to be addressed in future studies, our data strongly suggest that chromatin in MN is not a potent activator of cGAS-STING pathway, and that cGAS accumulation in MN is not a reliable marker of its activation.

The idea that chromatin is inert to cGAS even in the cytosol is also supported by the absence of cGAS activation by confinement-induced PN envelope rupture (Gentili et al, 2019). Instead, our data from irradiated cells suggest that cGAS is activated independently of MN. We do not exclude the possibility that small chromatin fragments that cannot be detected by mCitrine-NLS nor the cGAS reporter were leaked into the cytosol and became the source of cGAS-activating DNA. However, the complete absence of the interferon response in SCF-induced FuVis2 cells, which potentially harbor acentric X chromosome fragments in the cytosol, argues against this possibility. Instead, cumulative evidence supports the notion that nucleotides from disrupted mitochondria trigger the cGAS response in irradiated cells (Tigano et al, 2021; Guan et al, 2023). It is conceivable that cytosolic chromatin fragments rather inhibit cGAS activation in the presence of mtDNA.

Cytoplasmic chromatin fragments have been linked to inflammation and antitumor mechanisms due to their cGAS-accumulating potency (Dou et al, 2017; Glück et al, 2017; Harding et al, 2017; Mackenzie et al, 2017; Yang et al, 2017). However, our results argue against this hypothesis and suggest that MN is inert to cGAS-dependent innate immune pathway, raising the possibility that MN is more prone to developing chromosome abnormalities, including chromothripsis (Zhang et al, 2015; Ly et al, 2016, 2019; Kneissig et al, 2019; Umbreit et al, 2020) and epigenetic abnormalities (Agustinus et al, 2023; MacDonald et al, 2023), even in cells with an intact cGAS-STING pathway. Although our current study is limited to the specific reporter system, cytoplasmic mtDNA release needs to be carefully considered for studying cGAS-dependent inflammatory responses in different cellular contexts with MN formation.

## MATERIALS AND METHODS

### Cell Culture

Human colon carcinoma HCT116 cells (ATCC: American Type Culture Collection) and their derivatives were cultured in Dulbecco’s Modified Eagle Medium (DMEM, Nissui Pharmaceutical) supplemented with 10% fetal bovine serum (FBS), 2 mM L-glutamine, 0.165% NaHCO_3_, 100 U/ml penicillin/streptomycin, and 5 μg/ml Plasmocin (InvivoGen) and maintained at 37 °C in 5% CO_2_. Where indicated, the medium was supplemented with compound 3 (Selleckchem) and doxycycline (Sigma-Aldrich). For SPY650 (Cytoskeleton), cells were incubated with SPY650-containing medium for 2 hours before live-cell imaging, following a 5% concentration of the manufacturer’s instruction. During SGC0946 treatment, the medium containing SGC0946 was replaced every day for 3 days, and cells were transduced with the virus encoding Cas9-sgF21 in the presence of SGC0946 until and during live-cell imaging analysis.

### Plasmids

All plasmids used in this study are listed in Table S1. For cloning of the Sister-Control (SC) cassette plasmid (pMTH857) used for genomic integration, synthetic DNA fragments (Integrated DNA Technologies) were introduced into the original sister cassette plasmid (pMTH397)(Kagaya et al, 2020). A loxP sequence and two roxP sequences were inserted in downstream of a 5’ exon of mCitrine/mCerulean3 and neoR-franking regions, respectively, for potential future experiments. LentiCRISPR.v2 (addgene #52961) was mutagenized to introduce R691A to generate HiFi Cas9. pCAG-enAsCas12a-HF1(E174R/N282A/S542R/K548R)-NLS(nuc)-3xHA (addgene #107942) was used to obtain Lenti-enAsCas12a-HF1-2C-NLS, during which one more NLS was added to the C-terminus to improve its efficiency (Liu et al, 2019). LentiGuide-puro (addgene #52963) and an improved sgRNA scaffold sequence from pKLV2-U6gRNA5(Empty)-PGKBFP2AGFP-W (addgene #67979) were used to generate LentiGuide-puro-sgFUSION21-C+5bp plasmid. pH2B-miRFP703 (addgene #80001) and pCSII-EF-mVenus-hGeminin(1/110) (RDB15271) were used to generate pCSII-EF-emiRFP703-Geminin(1-110), during which the N-terminal sequence of miRFP703 was modified to obtain emiRFP703 (Matlashov et al, 2020). An improved rtTA3G was artificially synthesized (Integrated DNA Technologies) to obtain pLenti-rtTA3G (Zhou et al, 2006). pCW-Cas9 (addgene #50661) was modified to generate pTRE3G-miRFP670nano-p2a-Cas9(HiFi), during which the puroR-t2a-rtTA sequence was removed. miRFP670nano was artificially synthesized (Integrated DNA Technologies) (Oliinyk et al, 2019). Mutagenesis on Cas9 and cGAS was performed by conventional PCR followed by HiFi DNA Assembly (NEB) or In-Fusion cloning (Takara Bio). Full-length sequences of plasmids used in this study will become available at a public data share server upon publication.

### CRISPR/Cas9-mediated homology-directed DNA cassette integration into genomic DNA

The SC cassette (pMTH857) was integrated into the subtelomere locus on the short arm of the X chromosome in HCT116 cells, as described previously (Kagaya et al, 2020). Details of the validation steps are provided in the Supplementary information.

### Virus transduction

Lentivirus particles were generated as previously described (Kagaya et al, 2020) with minor modifications. Briefly, 1.6 µg of a transfer plasmid was transfected into LentiX 293T cells (Clontech Laboratories, inc.) with 0.8 µg of psPAX2 (addgene #12260) and 0.8 µg of pCMV-VSV-G (addgene #8454) using 9.6 µl of 1 mg/ml polyethylenimine (PEI) in a 6-well plate. The medium was replaced on the next day, and the medium containing lentivirus particles was collected on days 2 and 3 post-transfection and filtered through a 0.45 µm PES syringe filter (TELS25045, technolabsc inc.). For lentivirus infection, the medium of target cells was replaced with virus-containing medium supplemented with 8 µg/ml polybrene. Viral titers required for near 100% transduction were empirically determined by serial dilution of the virus-containing medium, followed by antibiotic selection. For the generation of cGAS and STING reporter-expressing cells, transduced cells were sorted by a fluorescence-activated cell sorter SH800S (Sony) with 130 µm sorting chips (Sony). For LentiCRISPR(HiFi) (Vakulskas et al, 2018), Lenti-enAsCas12a-HF1-2C (Kleinstiver et al, 2019), LentiGuide-sgRNA and pLKO.1-shRNA, transduced cells were selected by 1 µg/ml puromycin for 2 days after day 2 of transduction. The following guide sequences and shRNA sequences were used (5’ to 3’): sgFusion11, GTAGCGAACGTGTCCGGCGT; sgFusion21, ATTCTACCACGGCAGTCGTT; sgFusion22, GAACGTTGGCACTACTTCAC; sgFusion23, GTGGTAGAATAACGTATTAC; sgFusion24, GGATCCGTAGCGAACGTGTC; sgFusion25, AACGCCGGACACGTTCGCTA; sgFusion26, CGTTCCGGTCACTCCAACGC; crFusion6, AATAATGCCAATTATTTAAA; crFusion7, AATAATTGGCATTATTTAAA; crFusion8, AATAATGCCAATTATTTAAA; crFusion9, AGAAAAGCGATTTGGATTA; crFusion10, GATTATAACTTCGTATAGCA; crFusion11, AAGTTAAATTCATAACTTCG; crFusion12, ACTTTAAATAATGCCAATTA; crFusion13, ACTTTAAATAATTGGCATTA; crFusion14, AAGTTAAATTCACTCCAGA; shScramble, CCTAAGGTTAAGTCGCCCTCG; shcGAS, TTAGTTTTAAACAATCTTTCCT. For the generation of dox-inducible Cas9 (iCas9) cells, XpSC33 cells were simultaneously transduced with viruses encoding rtTA3G (pMTH1190) and TRE-promoter-driven miRFP670nano-p2a-Cas9(HiFi) (pMTH1197), exposed to 1 µg/ml doxycycline at 2 days post-transduction for 2 days, and sorted for miRFP670nano expression by the SH800S sorter with 130 µm sorting chips. Details of the validation for the iCas9 cell clones are found in the Supplementary information. For the generation of emiRFP703-Geminin-expressing cells, XpSC33-iCas9-20 cells were transduced with lentivirus encoding emiRFP703-Geminin (pMTH1094), a derivative of the FUCCI reporter for visualization of cells in S/G2/M phases of the cell cycle (Sakaue-Sawano et al, 2008). Then, emiRFP703-positive, and –negative cells were sequentially sorted by the SH800S sorter with 11 days intervals to enrich cells properly expressing the Geminin reporter. For the irradiation experiment, cells were transduced with lentivirus encoding mCitrine-NLS (pMTH1527) 4 days prior to irradiation.

### Flow Cytometry

Cells were collected by trypsinization, resuspended in cold 1x PBS containing 2.5 mM EDTA, and filtered through a 5 ml polystyrene round-bottom tube with a cell-strainer cap (Corning). Cells were analyzed using the SH800S cell sorter with 100 µm or 130 µm sorting chips (Sony). Single cells were gated based on their low FSC-W value before analysis and sorting. Fluorescence signals were detected using the following laser and filter combinations: DAPI and BFP, 405 nm laser, 450/50 filter; mCerulean3, 488 nm laser, 450/50 filter; GFP and mCitrine, 488 nm laser, 525/50 filter; mScarlet and mRuby3, 561 nm laser, 600/60 filter; and miRFP670nano and emiRFP703, 638 nm laser, 665/30 filter.

### Gamma ray irradiation

Two days prior to gamma-ray irradiation, cells were seeded onto a 35-mm dish. Subsequently, the cells were exposed to 1 Gy of gamma-rays using the Cs-137 Gammacell 40 Exactor (Best Theratronics Ltd.). Following irradiation, live-cell imaging was promptly carried out on the irradiated cells.

### Live-cell imaging

For the FuVis2 reporter experiment, XpSC33 and its derivative clones were transduced with lentivirus encoding Cas9-sgF21 or sgF21 (for iCas9-20 cells). Subsequently, these cells were seeded onto conventional cell culture dishes or plates at 2 days post-infection and subjected to live-cell imaging at 4 days post-infection. Live-cell imaging was performed as previously described (Kagaya et al, 2020). Briefly, cell culture dishes or plates were positioned on the BZ-X710 fluorescence microscope (KEYENCE), which was equipped with a metal halide lamp, stage-top chamber, and temperature controller featuring a built-in CO_2_ gas mixer (INUG2-KIW; Tokai hit). Each fluorescence signal was detected using the following filter cubes (M square): mCitrine, (ex: 500/20 nm, em: 535/30 nm, dichroic: 515LP); GFP, (ex: 470/40 nm, em: 525/50 nm, dichroic: 495LP); mScarlet and mRuby3, (ex: 545/25 nm, em: 605/70 nm, dichroic: 565LP); and emiRFP703 and SPY650, (ex: 620/60 nm, em: 700/75 nm, dichroic: 660LP). Images were captured using the BZ-H3XT time-lapse module, typically at intervals of 12 or 15 minutes, over a duration exceeding 60 hours. The formation of MN, the localization pattern of cGAS and the St-AI were analyzed through manual inspection.

### St-AI analysis

The cellular membrane of a target cell in the phase-contrast channel were manually inspected and tracked at 60-minute intervals using the freehand selection tool within the Fiji software (Schindelin et al, 2012). The tracked data was organized and stacked within the ROI (region of interest) manager. Subsequently, the stacked ROI data was superimposed onto the red channel (mRuby3-STING) to measure both maximum and mean intensities of mRuby3-STING within each cell lineage. For every cell and time-point, the maximum intensity of mRuby3-STING was divided by the mean intensity of mRuby3-STING, resulting in the computation of the STING Accumulation Index (St-AI).

### Micronuclei isolation

MN isolation was performed as previously described (Mohr et al, 2021) with minor modification. Briefly, XpSC33-iCas9-20 emiRFP703-Geminin cells were transduced with a virus encoding sgF21 and cultured in medium containing 1 µg/ml doxycycline for 8 days. The cells were subsequently sorted based on their emiRFP703-Geminin expression using the SH800S sorter (Sony) to enrich cells in S/G2/M phases of the cell cycle. After sorting, the cells were washed and then lysed using a lysis buffer [10mM pH 8.0 Tris-HCl, 2 mM Magnesium acetate, 3 mM Calcium chloride, 0.32 M Sucrose, 0.1 mM pH 8.0 EDTA, and 0.1% Nonidet P-40]. Putative MN and PN fractions were subsequently collected by sucrose gradient centrifugation. This process involved mixing 10 mL of the cell lysate with 15 ml of 1.6 M Sucrose buffer and 20 ml of 1.8 M Sucrose buffer, both containing 5 mM Magnesium Acetate and 0.1 mM pH 8.0 EDTA. The centrifugation was carried out at 950 x g for 20 minutes at 4 °C. The obtained putative PN and MN fractions were diluted with five times their volume in cold 1x PBS and centrifuged again at 950 x g for 20 minutes at 4 °C. After centrifugation, supernatants were discarded, and the pellet was resuspended in cold 1x PBS/0.1 mM EDTA with 0.1 µg/ml DAPI for subsequent sorting.

### Fluorescent In Situ Hybridization

For mitotic chromosome spread, XpSC33 Cas9-sgF21 cells were exposed to 100 ng/ml colcemid on day 6 post-infection for 16 hours to enrich mitotic cells. Subsequently, the cells were sorted based on mCitrine and mCerulean3 fluorescent cells using the SH800S sorter. The sorted cells were pelleted and then exposed to a 5 ml solution of 75 mM KCl for 7 minutes at room temperature. The swelling process was halted by adding 0.5 ml of ice-cold 3:1 Methanol/Acetic acid, and the cells were pelleted again for fixation in a 5 ml ice-cold 3:1 Methanol/Acetic acid solution. After centrifugation and resuspension in fresh ice-cold 3:1 Methanol/Acetic acid, the cells were deposited onto glass slides. Following air drying, the cells were mounted with an XCP X orange probe specific for the entire X chromosome (MetaSystems Probes) and an XCE X/Y green/orange probe for X/Y chromosome centromeres (MetaSystems Probes), following the manufacturer’s instructions. For samples enriched with MN and PN, sorted samples were centrifuged at 950 x g for 20 minutes at 4 °C to eliminate the supernatant. The pellets were then resuspended in 150 µl of ice-cold 3:1 methanol/acetic acid, and the samples were deposited onto glass slides. After air drying, the samples were mounted with the XCP X orange probe (MetaSystems Probes) beneath coverslips, heated at 75 °C for 2 minutes, and incubated at 37 °C for overnight. Slides were subjected to washing with 0.4 x SSC at 72 °C for 2 minutes and 2 x SSC with 0.05% Tween-20 at room temperature for 30 seconds, followed by rinsing with distilled water. After a brief drying period, samples were mounted using PNG anti-fade [4% n-propyl gallate, 100 mM Tris pH8.5, 90% glycerol] with 0.1 µg/ml DAPI.

### Immunofluorescence

Cells were cultured on coverslips coated with Alcian Blue 8GX (A5268, Sigma-Aldrich), fixed with 4% paraformaldehyde in 1x PBS for 15 minutes at room temperature, and washed with 1x PBS three times. The fixed cells were permeabilized using 0.2% Triton X-100, 0.02% Skim milk (nacalai tesque), and 0.02% BSA (Sigma-Aldrich) in 1x PBS for 5 minutes at room temperature in dark. After rinsing with 1x PBS once and then with PBST (0.1% Tween20, 1x PBS), the cells were incubated with Phospho-STING (Ser366) antibody (19781S, Cell Signaling Technology) at a 1:200 dilution in PBST for 45 minutes at room temperature. Following three washes with PBST, the cells were incubated with Alexa-488-conjugated anti-rabbit (A11034, Invitrogen) or Alexa-594-conjugated anti-rabbit (ab150080, abcam) at a 1:1000 dilution in PBST for 45 minutes at room temperature in dark, and then washed with PBST and milliQ water. After air drying, coverslips were mounted on glass slides using PNG anti-fade.

### RT-qPCR

Total RNA was extracted from cells by the RNeasy Mini Kit (Qiagen). Then, 0.165 µg of total RNA was reverse transcribed using 62.5 nM Oligo dT and 0.18 µl of AMV reverse transcriptase (NIPPON GENE) in a total 25 µl reaction mix following the manufacturer’s instructions. The resulting cDNA was used for qPCR with THUNDERBIRD Next SYBR pRCR Mix (Toyobo) and the StepOnePlus Real-Time PCR System (Applied Biosystems).

### Statistical analysis

All statistical analyses and graphing were performed using GraphPad Prism software (version 10.0). For numerical data with assumed Gaussian distribution, two-tailed Student’s t-test and ordinary one-way ANOVA followed by post-hoc comparisons were applied. For categorical data, the chi-square test was utilized. The alpha level was set at 0.05.

## ACKNOWLEDGEMENTS

We thank the Drug Discovery Centre, supported by the iSAL (innovative support alliance for life science) Kyoto University for the cell sorter; CORE Program of the Radiation Biology Center, Kyoto University for gamma ray irradiation; the RIKEN BRC through the National BioResource Project of the MEXT, Japan for material distribution; Dorus Gadella, Fuyuki Ishikawa, Keith Joung, Masato Kanemaki, Jan Karlseder, Benjamin Kleinstiver, Eric Lander, Atsushi Miyawaki, David Sabatini, Ryota Sato, Tomohiko Taguchi, Didier Trono, Vladislav Verkhusha, Robert Weinberg, Kousuke Yusa and Feng Zhang for sharing materials; Yumi Hayashi for assistance with molecular cloning and St-AI analysis; Andrea Ruelas-Gonzalez and Yuya Nishida for assistance with St-AI analysis; and members of the Hayashi lab for their suggestion and discussion. This project was supported by grants from the Grant-in-Aid for Scientific Research (B) (20H03183) to M.T.H.; and institutional fundings from the Hakubi Center and the Graduate School of Medicine, Kyoto University to M.T.H.

## AUTHOR CONTRIBUTIONS

Y.S. and M.T.H. conceived the study. M.T.H. cloned the FuVis2 reporter cell. Y.S. performed experiments and data analysis. M.T.H. assisted with data analysis. M.T.H. secured funding. Y.S. and M.T.H. wrote the manuscript.

## DATA AVAILABILITY

All data are archived at Kyoto University and available from the corresponding author upon reasonable request. Full-length DNA sequences of plasmids used in this study will become available at a public data share server upon publication.

## ADDITIONAL INFORMATION

Supplementary information is available.

## Figure legends

**Figure S1.**
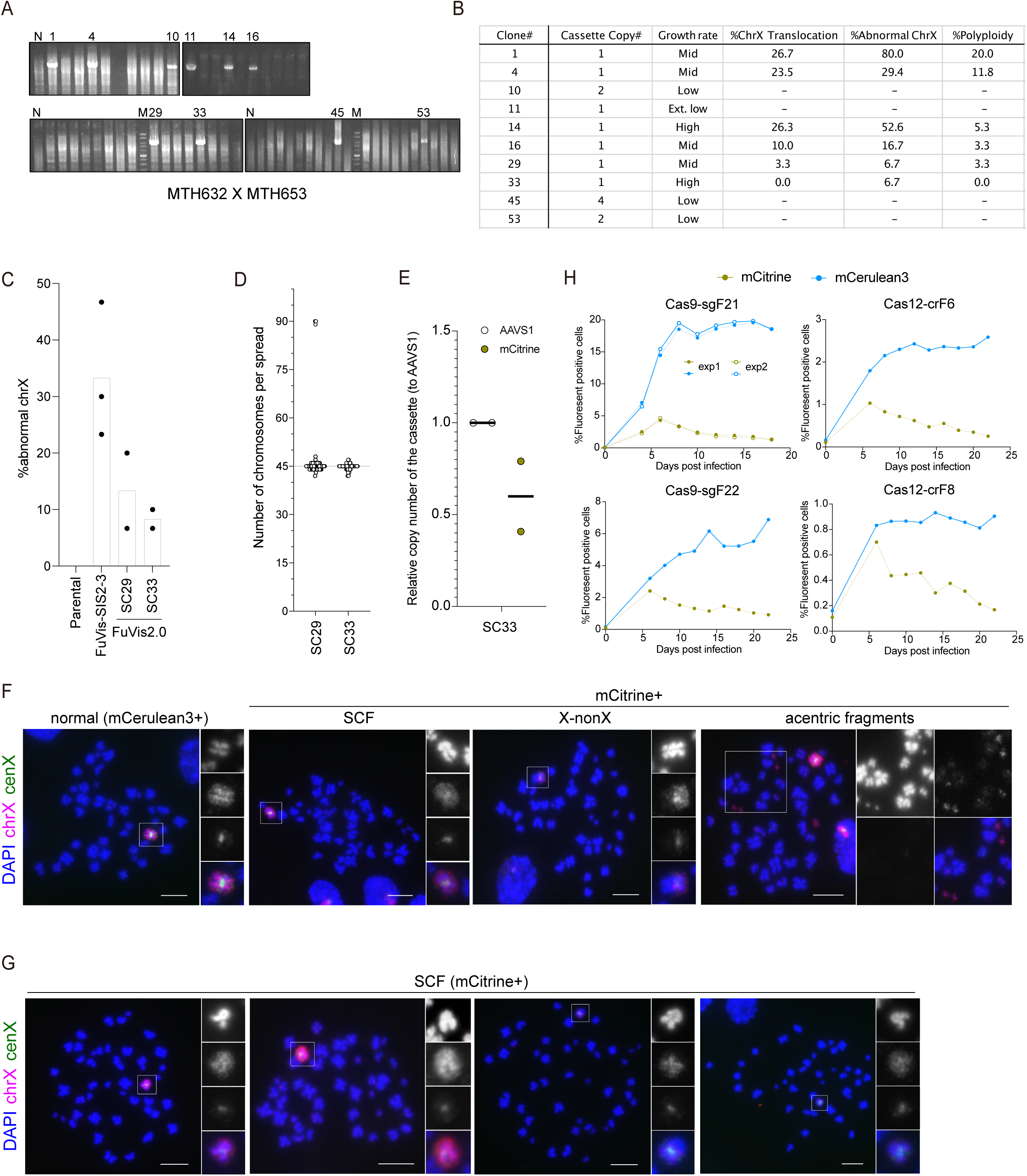
Development and validation of FuVis2-XpSC cells. (A) Agarose gel images of genomic PCR products (5,525 bp) from FuVis2-XpSC candidate clones. The primers used are indicated in Table S2. (B) Table presenting characteristics of each candidate clone selected from the genomic PCR in (A). The cassette copy number was determined through quantitative genomic PCR. Growth rates were examined by culture passage rate. X Chromosome abnormalities were analyzed by FISH using whole chrX and cenX probes. (C) Percentage of cells exhibiting abnormal chrX. FuVis-SIS2-3 represents the first generation FuVis reporter. Cells were cytologically analyzed by FISH using whole chrX and cenX probes (n = 30/experiment, three biological replicates for FuVis-SIS2-3, two biological replicates for FuVis2-XpSC29 and XpSC33). (D) Number of total chromosomes per metaphase spread in the indicated candidates. (E) Relative copy number of the integrated reporter cassette in FuVis2-XpSC33 clone. A plasmid carrying one copy of both AAVS1 and mCitrine sequences was used as a standard in quantitative PCR analysis. The AAVS1 locus on the genome (2 copies) was used as a control. (F) Representative images of the X chromosome in XpSC33 Cas9-sgF21 cells as shown in Fig. 1C. Whole metaphase spreads corresponding to Fig. 1C are shown. Scale bar, 10 µm. (G) Representative images of SCF on the X chromosome in XpSC33 Cas9-sgF21 cells as shown in Fig. 1C. Scale bar, 10 µm. (H) Time-course of the percentage of mCerulean3– and mCitrine-positive cells in XpSC33 cells expressing indicated endonucleases and guide RNAs. Cells were transduced with viruses encoding endonucleases and guide RNA, selected by puromycin from 2 days post-infection, and analyzed by flow cytometry.

**Figure S2.**
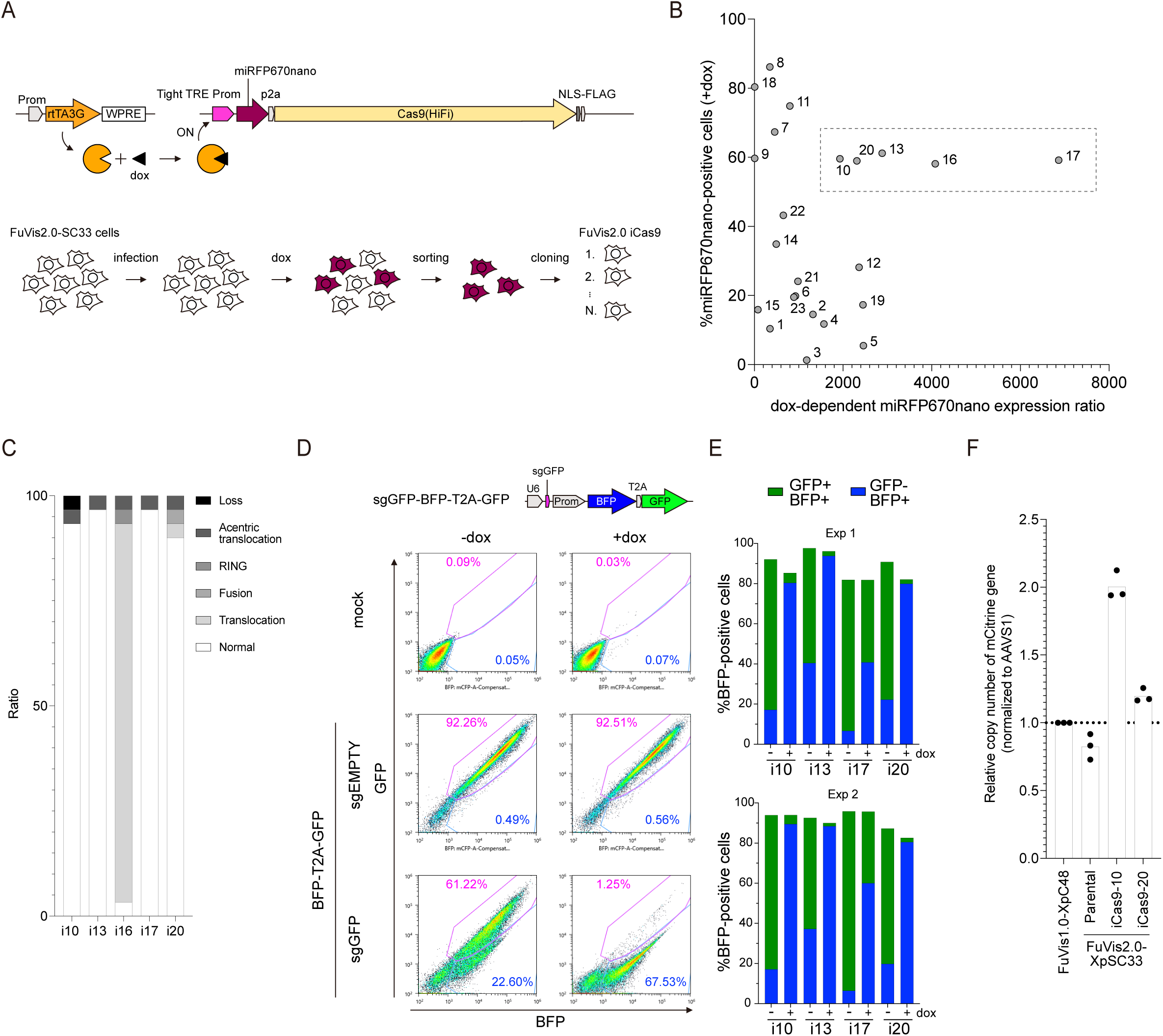
Development and validation of FuVis2-XpSC33 iCas9 reporter cells. (A) Schematic of the doxycycline (dox)-inducible Cas9(HiFi) system. SC33 cells were transduced with two viruses encoding rtTA3G and a tight TRE promoter-regulated miRFP670nano-p2a-Cas9(HiFi). The transduced cells were treated with 1 µg/ml dox for 2 days, and miRFP670nano-positive cells were sorted to obtain subclones. (B) Scatter plot displaying the percentage of miRFP670nano-positive cells and dox-dependent miRFP670nano expression ratio in individual subclones from (A). Subclones were grown with or without 1 µg/ml dox for 2 days and analyzed by flow cytometry. The dox-dependent expression ratio was calculated by dividing the percentage of miRFP670nano-positive cells in the presence of dox by the percentage in the absence of dox. The dot line indicates subclones selected for subsequent validation. (C) Percentage of structural and numerical abnormalities of chrX in the indicated subclones. Metaphase chromosome spreads were analyzed by FISH using whole chrX and cenX probes. Abnormalities include loss (loss of the entire chrX), acentric translocation (acentric fragment of chrX translocated to another chromosome), RING (fusion between the long-arm and the short-arm telomeres of chrX), fusion (fusion between chrX and another chromosome), and translocation (non-X chromosome fragment translocated to chrX). (D) Evaluation of the dox-inducible Cas9(HiFi) efficiency in the subclones. Subclones were transduced with a virus encoding a Cas9 reporter cassette (top) and exposed to 0.1 µg/ml dox for 1 day, then harvested at 4 days post-infection for FACS analysis. GFP and BFP intensities are plotted (bottom). Results from iCas9-10 are shown as representative. Blue gates (BFP-positive and GFP-negative population) and magenta gates (GFP– and BFP-positive population) indicate cells that possess the reporter with and without, respectively, Cas9-dependent indel on the GFP gene. Percentages of total single cells are shown. (E) Percentage of BFP-positive cells with or without GFP expression in indicated subclones expressing sgGFP as analyzed in (D). Results from two biologically independent experiments are shown. (F) Relative copy number of the integrated SC reporter cassette in the indicated subclones. A plasmid carrying one copy of both AAVS1 and mCitrine sequences (pMTH864) was used as a standard in quantitative PCR analysis. The AAVS1 locus on the genome (2 copies) was used to normalize the copy number. FuVis1-XpC48 serves as a control that carries a single copy of the cassette (mCitrine gene).

**Figure S3.**
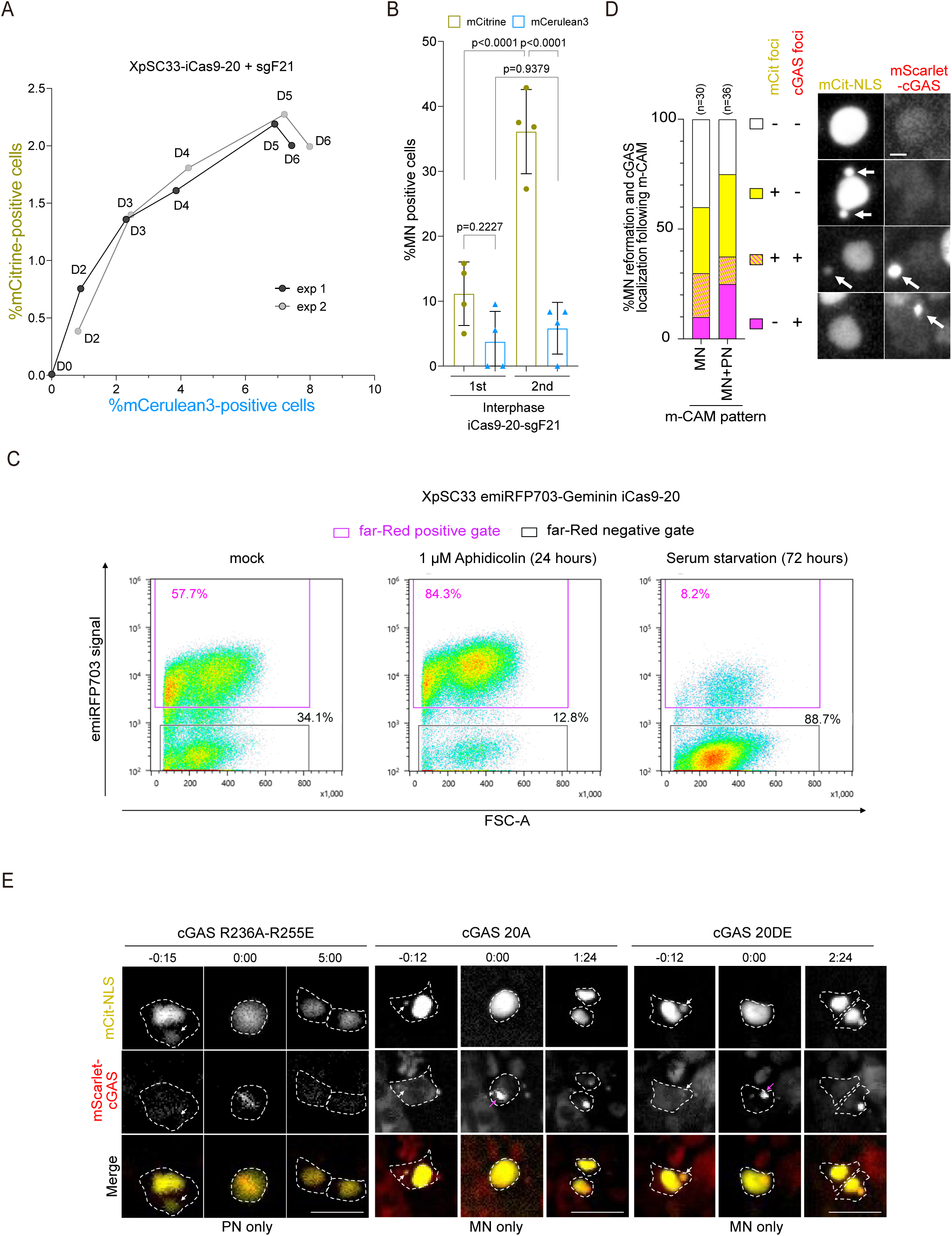
Micronuclei derived from sister chromatid fusion are captured by cGAS during and after mitosis. (A) Time-course of the percentage of mCerulean3– and mCitrine-positive cells in XpSC33-iCas9-20 sgF21 cells. Cells were transduced with a virus encoding sgF21 and exposed to 0.1 µg/ml dox for one day. Cells were analyzed every day until 6 days post-infection. (B) Percentage of MN-positive cells in XpSC33-iCas9-20 sgF21 cells at the indicated count of the cell cycle. Cells were subjected to live-cell imaging at 4 days post-infection. Results from four biological replicates are shown (n = at least 18/experiment, ordinary one-way ANOVA). (C) Flow cytometry analysis of emiRFP703-Geminin expression in XpSC33 iCas9-20 cells. Cells were either treated with 1 µM aphidicolin for 24 hours or serum-free medium for 72 hours before harvesting. Percentages of emiRFP703-positive cells are shown. (D) Percentage of cells with indicated mCitrine-NLS and mScarlet-cGAS localization patterns after the m-CAM event (left). XpSC33 mScarlet-cGAS Cas9-sgF21 cells were subjected to live-cell imaging at 4 days post-infection. Representative images of each localization pattern are shown (right). White arrows represent mCitrine-NLS or mScarlet-cGAS foci that implicate MN formation. Scale bar, 10 µm. (E) Representative live-cell images of mScarlet-cGAS localization upon NEBD (0:00) in MN-positive XpSC33 Cas9-sgF21 cells expressing the indicated mScarlet-cGAS mutants. NEBD is recognized by the diffusion of mCitrine-NLS signal. White arrows indicate MN with intact membrane before NEBD; magenta arrows indicate mScarlet-cGAS foci on MN-derived chromatin upon NEBD. Dot lines indicate the cellular membrane determined by phase-contrast images. cGAS localization pattern is indicated below for each condition. Scale bar, 25 µm.

**Figure S4.**
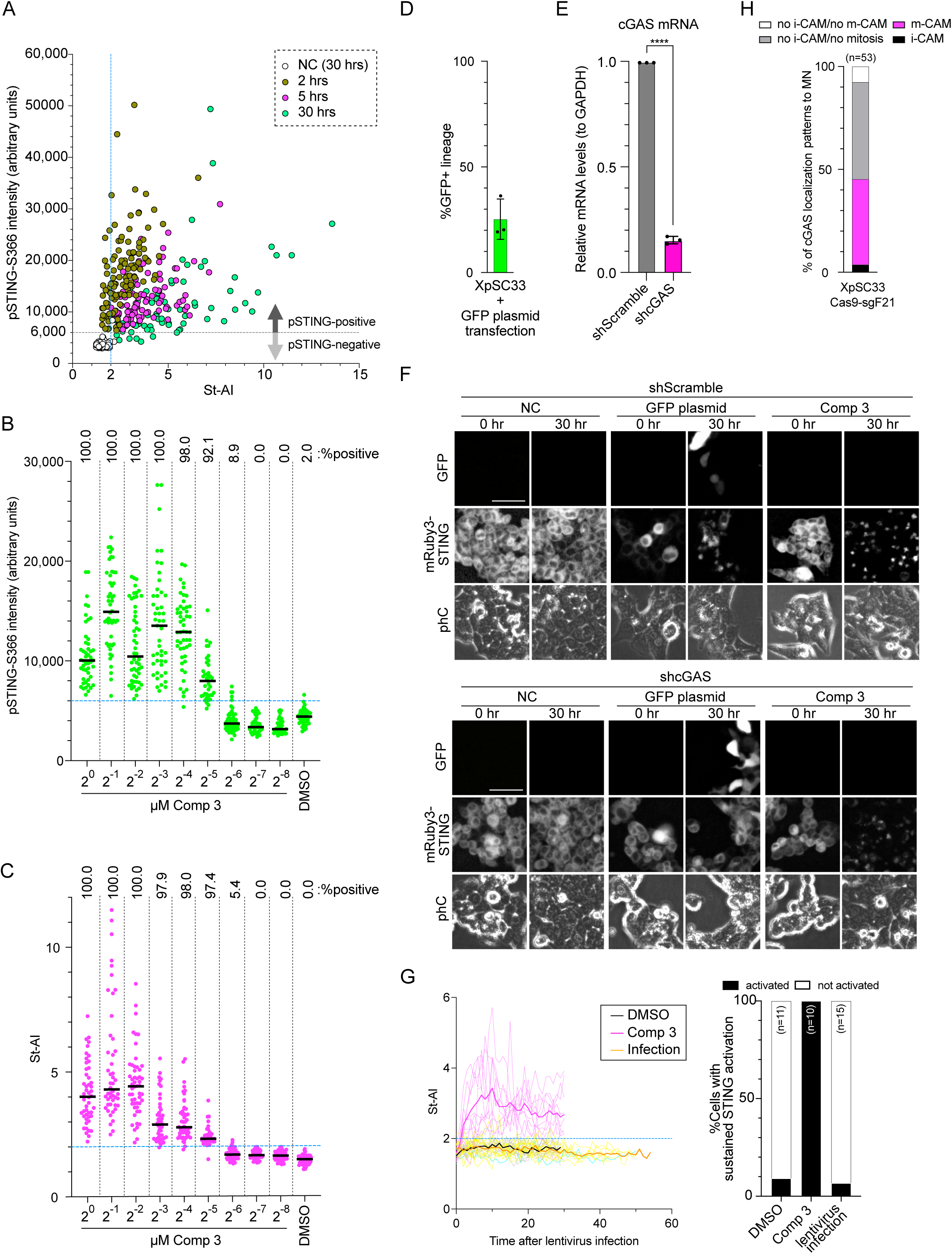
Validation of mRuby3-STING reporter. (A) Scatter plot of pSTING-S366 intensity and St-AI in XpSC33 emiRFP703-cGAS mRuby3-STING cells exposed to 1 µM compound 3 for indicated hours. The gray dot line indicates a threshold of pSTING-S366 intensity (6,000), above which signal was defined as positive. The blue dot line indicates a threshold of St-AI (2.0), above which STING was defined as accumulated and active. (B) pSTING-S366 signal intensity in XpSC33 emiRFP703-cGAS mRuby3-STING cells exposed to the indicated dose of compound 3 for 5 hours. Percentages of pSTING-positive cells is indicated above. (C) St-AI in the same sample from (B). Percentages of cells with St-AI more than 2 are indicated above. (D) Percentage of GFP-positive lineages in XpSC33 emiRFP703-cGAS mRuby3-STING cells transfected with pMAX-TurboGFP plasmid. Individual cells at transfection were tracked by live-cell imaging to determine whether their descendants express GPF or not. Three biological replicates are shown (n = at least 30 lineages/experiment, mean +/− s.d.). (E) Relative mRNA level of cGAS normalized to GAPDH. XpSC33 emiRFP703-cGAS mRuby3-STING Cas9-sgF21 cells were transduced with shScramble or shcGAS viruses and selected by puromycin for 2 days from 2 days post-infection. Total RNA was extracted and subjected to RT-qPCR (n = 3 biological replicates, unpaired t-test). ***p<0.0001. (F) Representative images of GFP and mRuby3-STING at 30 hours after plasmid transfection or compound 3 treatment. XpSC33 emiRFP703-cGAS mRuby3-STING cells transduced with viruses encoding indicated shRNA were transfected with pMAX-TurboGFP or exposed to 1 µM compound 3 at 4 days post-infection. Scale bar, 50 µm. (G) (Left) Time-course analysis of St-AI after 1 µM compound 3 treatment or virus transduction of XpSC33 mRuby3-STING cells. Cells were transduced with a virus encoding miRFP670, and only miRFP670-positive cells were analyzed after infection. Bold lines indicate the average of individual cells in each condition. The blue dot line represents St-AI = 2.0. (Right) Percentages of cells exhibiting sustained STING activation (St-AI > 2.0 for a duration over 4 hours) in each condition. (H) Percentage of cGAS localization patterns to MN in XpSC33 emiRFP703-cGAS mRuby3-STING Cas9-sgF21 cells. MN-positive cells were tracked for emiRFP703-cGAS accumulation. No i-CAM/no m-CAM indicates MN-positive cells entered mitosis without any signs of m-CAM.

**Figure S5.**
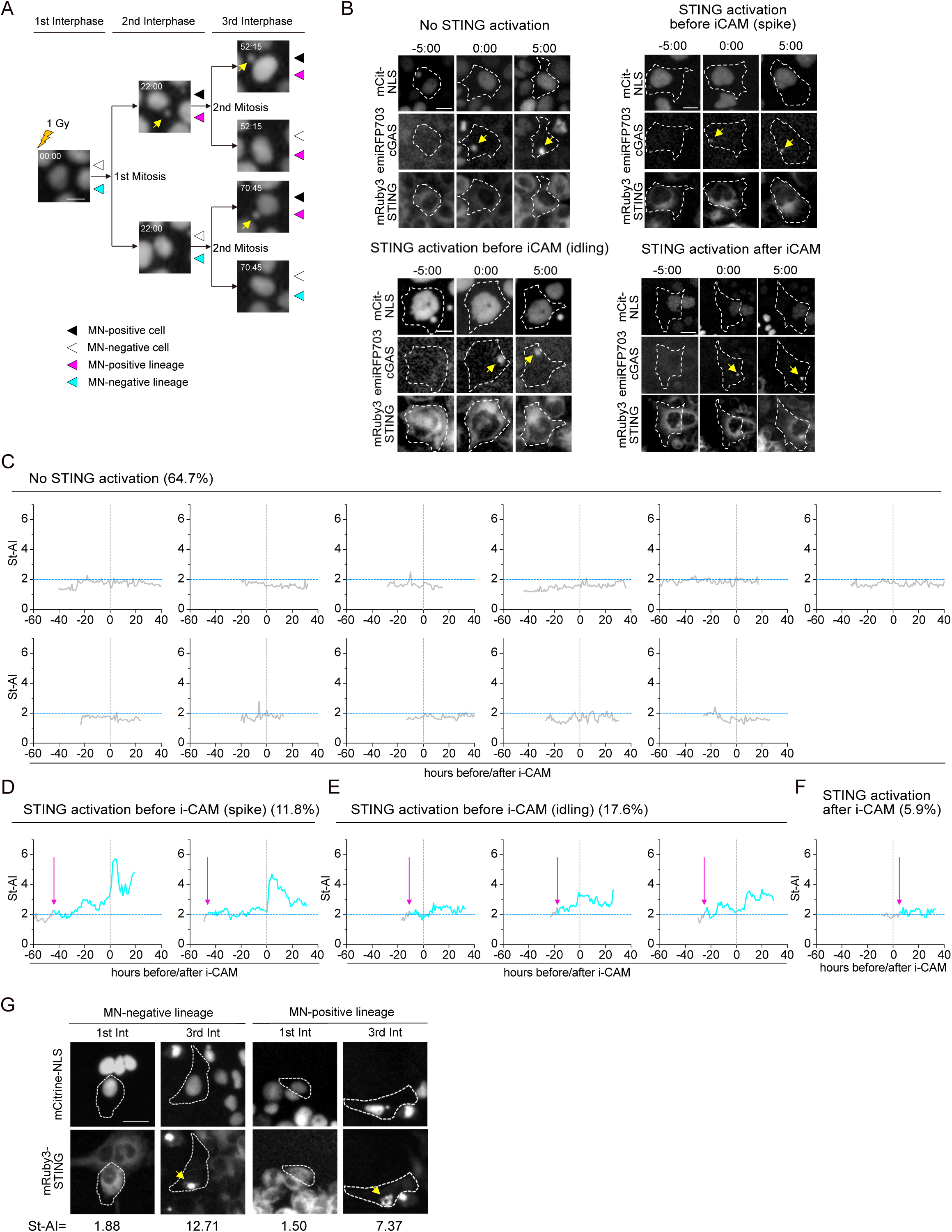
The i-CAM event after irradiation is not sufficient to activate STING. (A) Representative images of XpSC33 mCitrine-NLS emiRFP703-cGAS mRuby3-STING cells in each cell cycle count after irradiation as shown in Fig. 5A. Only mCitrine signal is shown. Yellow arrows indicate MN. Scale bar, 25 µm. (B) Representative images of irradiated cells 5 hours before and after i-CAM (0:00) analyzed as in (A). Yellow arrows indicate the i-CAM event. White dot lines represent cellular membranes determined by phase-contrast images. Scale bar, 25 µm. (C-F) Kinetics of St-AI in cells that showed i-CAM after 1 Gy IR. XpSC33 mCitrine-NLS emiRFP703-cGAS mRuby3-STING cells were exposed to 1 Gy IR and subjected to live-cell imaging. Vertical gray dot lines indicate the time at the i-CAM event. Horizontal blue dot lines indicate St-AI = 2.0. Magenta arrows indicate the time at sustained STING activation started. Names and percentages of each category of STING activity are shown above. (G) Representative images of XpSC33 mCitrine-NLS emiRFP703-cGAS mRuby3-STING cells after 1 Gy IR in MN-negative and MN-positive lineages. Yellow arrows indicate STING accumulation. St-AI is indicated below for each condition. Scale bar, 25 µm.

## Supplementary information

### Establishment and validation of FuVis2-XpSC cell clones

The Sister-Control (SC) reporter cassette (pMTH857) was integrated into a telomere-adjacent subtelomere sequence on the short arm of the X chromosome in HCT116 cells through CRISPR/Cas9-directed homology-mediated recombination (HR) facilitated by pMTH393 (Fig. 1A). We successfully isolated 56 independent G418-resistant clones during this process. Subsequently, we validated ten clones (SC1, SC4, SC10, SC11, SC14, SC16, SC29, SC33, SC45, SC53) for their intended integration using genomic PCR (Fig. S1A). Quantitative PCR analysis of the integrated reporter cassette revealed that three clones (SC10, SC45, SC53) carried two or more copies of the integrated reporter (Fig. S1B). Besides these three clones, one clone (SC11) displaying an exceptionally low growth rate was excluded from the pool of candidate clones (Fig. S1B). Further examination of the X chromosome structure in the remaining six candidate clones was conducted through FISH analysis using DNA probes spanning whole X chromosome (chrX) and the X centromere (cenX). The results indicated that two clones (SC29, SC33) harbored relatively normal X chromosome (Fig. S1B, C). Since the clone SC29 exhibited tetraploidy within the population (Fig. S1D), we chose the clone SC33 for subsequent analysis.

### Establishment and validation of FuVis2-XpSC33-iCas9 cell clones

To establish XpSC33 cells featuring doxycycline (dox)-inducible HiFi SpCas9, the cells were transduced with two independent viruses carrying rtTA3G under a constitutive promoter (pMTH1190) and miRFP670nano-p2a-Cas9(HiFi) under the tight TRE promoter (pMTH1197), respectively (Fig. S2A). The infected cells were treated with 1 µg/ml dox for two days and miRFP670nano-positive cells were sorted using the SH800S cell sorter, which was followed by single-cell subcloning (Fig. S2A). Resulting 23 subclones were subjected to a two-day dox treatment and subsequent FACS analysis to confirm the dox-dependent miRFP670nano expression (Fig. S2B). We identified five candidate subclones (iCas9-10, iCas9-13, iCas9-16, iCas9-17, iCas9-20), which displayed more than 50% miRFP670nano-positive cells and exhibited a substantial increase of more than 1,000 times in miRFP670nano-positive cells upon dox treatment (Fig. S2B). FISH analysis using chrX and cenX probes revealed that iCas9-16 harbored a translocation on the X chromosome (Fig. S2C). To assess Cas9 efficiency, we transduced the candidate subclones with a virus carrying a Cas9 reporter sequence (Fig. S2D). This analysis revealed that iCas9-10 and iCas9-20 displayed efficient GFP targeting activities upon dox exposure, with minimal background activities (Fig. S2D). Inspection of the copy numbers of the SC reporter cassette revealed that iCas9-10 carried a duplication of the SC reporter cassette (Fig. S2E). Given these assessments, we have selected XpSC33 iCas9-20 subclone for subsequent analysis.

