## Supplementary material for "Micronucleus Is Not a Potent Inducer of cGAS-STING Pathway": Table S1

**Table S1. Plasmids used in this study**

| pMTH_No | Name | Purpose | Source |
|---|---|---|---|
| 127 | pLKO.1-shScramble | shRNA control | Addgene 1864 |
| 380 | pMax-TurboGFP | DNA transfection control | Lonza Nucleofector kit |
| 393 | eSpCas9(1.1)-sgCHRXpYp-subtel2 | FuVis2-XpSC cloning | Kagaya et al. 2020 |
| 479 | LentiCRISPR(1.1)-sgFUSION11 | Cas9.v2 sgFUSION expression | Kagaya et al. 2020 |
| 857 | pXp-tr-sis-ctrl(neo)-rox-sg-mCer3C | FuVis2-XpSC cloning | This study |
| 864 | pBS-sister(neo)-AAVS1 | qPCR standard | Kagaya et al. 2020 |
| 907 | pLenti-bla-p2a-mScarlet-3FL-hcGAS | cGAS reporter | This study |
| 1001 | pLentiCRISPR(HiFi)-sgFusion21 | Cas9(HiFi) sgFUSION expression | This study |
| 1002 | pLentiCRISPR(HiFi)-sgFusion22 | Cas9(HiFi) sgFUSION expression | This study |
| 1003 | pLentiCRISPR(HiFi)-sgFusion23 | Cas9(HiFi) sgFUSION expression | This study |
| 1004 | pLentiCRISPR(HiFi)-sgFusion24 | Cas9(HiFi) sgFUSION expression | This study |
| 1005 | pLentiCRISPR(HiFi)-sgFusion25 | Cas9(HiFi) sgFUSION expression | This study |
| 1006 | pLentiCRISPR(HiFi)-sgFusion26 | Cas9(HiFi) sgFUSION expression | This study |
| 1011 | Lenti-enAsCas12a-HF1-2C-NLS-crFUSION6 | Cas12a-HF1 crFUSION expression | This study |
| 1012 | Lenti-enAsCas12a-HF1-2C-NLS-crFUSION7 | Cas12a-HF1 crFUSION expression | This study |
| 1013 | Lenti-enAsCas12a-HF1-2C-NLS-crFUSION8 | Cas12a-HF1 crFUSION expression | This study |
| 1014 | Lenti-enAsCas12a-HF1-2C-NLS-crFUSION9 | Cas12a-HF1 crFUSION expression | This study |
| 1015 | Lenti-enAsCas12a-HF1-2C-NLS-crFUSION10 | Cas12a-HF1 crFUSION expression | This study |
| 1016 | Lenti-enAsCas12a-HF1-2C-NLS-crFUSION11 | Cas12a-HF1 crFUSION expression | This study |
| 1017 | Lenti-enAsCas12a-HF1-2C-NLS-crFUSION12 | Cas12a-HF1 crFUSION expression | This study |
| 1018 | Lenti-enAsCas12a-HF1-2C-NLS-crFUSION13 | Cas12a-HF1 crFUSION expression | This study |
| 1019 | Lenti-enAsCas12a-HF1-2C-NLS-crFUSION14 | Cas12a-HF1 crFUSION expression | This study |
| 1050 | pLenti-bla-p2a-emiRFP703 | lentivirus infection control | This study |
| 1058 | pLenti-p2a-emiRFP703-3FL-hcGAS | cGAS reporter | This study |
| 1090 | pLenti-bla-p2a-mScarlet-3FL-hcGAS-R236A-R255E | cGAS reporter | This study |
| 1094 | pCSII-EF-emiRFP703-Geminin(1-110) | S/G2/M cell cycle reporter | This study |
| 1148 | pLenti-bla-p2a-mScarlet-3FL-hcGAS-20A | cGAS reporter | This study |
| 1149 | pLenti-bla-p2a-mScarlet-3FL-hcGAS-20DE | cGAS reporter | This study |
| 1159 | pMX-mRuby3-hSTING | STING reporter | This study |
| 1190 | pLenti-rtTA3G | dox-inducible Cas9(HiFi) | This study |
| 1197 | pTRE3G-miRFP670nano-p2a-Cas9(HiFi) | dox-inducible Cas9(HiFi) | This study |
| 1292 | pKLV2-U6gRNA5(Empty)-PGKBFP2AGFP-W | iCas9 validation | Addgene 67979 |
| 1293 | pKLV2-U6gRNA5(gGFP)-PGKBFP2AGFP-W | iCas9 validation | Addgene 67980 |
| 1366 | pLenti-p2a-emiRFP703-3FL-hcGAS-R236A-R255E | cGAS reporter | This study |
| 1367 | pLenti-p2a-emiRFP703-3FL-hcGAS-20A | cGAS reporter | This study |
| 1418 | LentiGuide-puro-sgFUSION21-C+5pb_stem | sgFUSION21 expression | This study |
| 1502 | pLKO.1-shcGAS | shRNA cGAS | This study |
| 1527 | pLenti-mCitrine-NLS | mCitrine-NLS expression | This study |

**Plasmids used for cloning of the plasmids used in this study**

| pMTH_No | Name | Purpose | Source |
|---|---|---|---|
| 221 | LentiCRISPR v2 | pLentiCRISPR(HiFi) sgFUSION construction | Addgene 52961 |
| 224 | LentiGuide-puro | LentiGuide-puro-sgFUSION21-C+5pb_stem construction | Addgene 52963 |
| 319 | pMXs-mCitrine | mCitrine-NLS construction | Lab stock |
| 420 | pLenti-blaR-p2a | pLenti-bla-p2a-emiRFP703 construction | Lab stock |
| 664 | pmScarlet_C1 | mScarlet-tag construction | Addgene 85042 |
| 751 | pH2B-miRFP703 | pLenti-bla-p2a-emiRFP703 construction | Addgene 80001 |
| 753 | pCW-Cas9 | pTRE3G-miRFP670nano-p2a-Cas9(HiFi) construction | Addgene 50661 |
| 776 | pLentiCRISPR(HiFi) | pLentiCRISPR(HiFi) sgFUSION construction | This study |
| 792 | pCAG-enASCas12a-HF1(E174R/N282A/S542R/K548R)-NLS(nuc)-3xHA | Cas12a-HF1 crFUSION construction | Addgene 107942 |
| 821 | Lenti-enAsCas12-HF1-2C-NLS | Cas12a-HF1 crFUSION construction | This study |
| 890 | pcDNA3-hcGAS-WT | cGAS reporter construction | Dr. Tomohiko Taguchi |
| 1031 | pMX-IP-mRuby3-mouseSTING | STING reporter construction | Dr. Tomohiko Taguchi |
| 1050 | pLenti-bla-p2a-emiRFP703 | pCSII-EF-emiRFP703-Geminin(1-110) construction | This study |
| 1080 | pMX4-puro-hSTING-FLAG-His | STING reporter construction | Dr. Ryota Sato |
| 1282 | pLenti-Clover-NLS | mCitrine-NLS construction | Lab stock |
| 1345 | LentiGuide-puro-sgFUSION21 | LentiGuide-puro-sgFUSION21-C+5pb_stem construction | This study |
