## Supplementary material for "Micronucleus Is Not a Potent Inducer of cGAS-STING Pathway": Table S2

**Table S2. Primers used in this study**

| MTH primer number | Name | 5'-sequence-3' | Purpose |
|---|---|---|---|
| 1546 | sgFusion21-fwd | caccgATTCTACCACGGCAGTCGTT | sgRNA |
| 1547 | sgFusion21-rev | aaacAACGACTGCCGTGGTAGAATc | sgRNA |
| 1548 | sgFusion22-fwd | caccgGAACGTTGGCACTACTTCAC | sgRNA |
| 1549 | sgFusion22-rev | aaacGTGAAGTAGTGCCAACGTTCc | sgRNA |
| 1550 | sgFusion23-fwd | caccgGTGGTAGAATAACGTATTAC | sgRNA |
| 1551 | sgFusion23-rev | aaacGTAATACGTTATTCTACCACc | sgRNA |
| 1552 | sgFusion24-fwd | caccgGGATCCGTAGCGAACGTGTC | sgRNA |
| 1553 | sgFusion24-rev | aaacGACACGTTCGCTACGGATCCc | sgRNA |
| 1554 | sgFusion25-fwd | caccgAACGCCGGACACGTTCGCTA | sgRNA |
| 1555 | sgFusion25-rev | aaacTAGCGAACGTGTCCGGCGTTc | sgRNA |
| 1556 | sgFusion26-fwd | caccgCGTTCCGGTCactccaACGC | sgRNA |
| 1557 | sgFusion26-rev | aaacGCGTtggagtGACCGGAACGc | sgRNA |
| 1578 | crFUSION6-fwd | AGATaataATGCCAATtatttaaaTTTTAT | crRNA |
| 1579 | crFUSION6-rev | AAAAATAAAAtttaaataATTGGCATtatt | crRNA |
| 1580 | crFUSION7-fwd | AGATaataATTGGCATtatttaaaTTTTAT | crRNA |
| 1581 | crFUSION7-rev | AAAAATAAAAtttaaataATGCCAATtatt | crRNA |
| 1595 | crFUSION8-fwd | AGATaataATGCCAATtatttaaaTTTTAT | crRNA |
| 1582 | crFUSION8-rev | AAAAATAAAAtttaaataATTGGCATtatt | crRNA |
| 1583 | crFUSION9-fwd | AGATagaaaagcgatttggattATTTTAT | crRNA |
| 1584 | crFUSION9-rev | AAAAATAAAATaatccaaatcgcttttct | crRNA |
| 1585 | crFUSION10-fwd | AGATgattATAACTTCGTATAGCATTTTAT | crRNA |
| 1586 | crFUSION10-rev | AAAAATAAAATGCTATACGAAGTTATaatc | crRNA |
| 1587 | crFUSION11-fwd | AGATaagttaaattcATAACTTCGTTTTAT | crRNA |
| 1588 | crFUSION11-rev | AAAAATAAAACGAAGTTATgaatttaactt | crRNA |
| 1589 | crFUSION12-fwd | AGATactttaaataATGCCAATtaTTTTAT | crRNA |
| 1590 | crFUSION12-rev | AAAAATAAAAtaATTGGCATtatttaaagt | crRNA |
| 1591 | crFUSION13-fwd | AGATactttaaataATTGGCATtaTTTTAT | crRNA |
| 1592 | crFUSION13-rev | AAAAATAAAAtaATGCCAATtatttaaagt | crRNA |
| 1593 | crFUSION14-fwd | AGATaagttaaattcactccaGATTTTAT | crRNA |
| 1594 | crFUSION14-rev | AAAAATAAAATCtggagtgaatttaactt | crRNA |
| 1833 | oligo-dT | TTTTTTTTTTTTTTT | cDNA (RT-PCR) |
| 2332 | cGAS-fwd | ATGCAAAGGAAGGAAATGGT | qPCR |
| 2333 | cGAS-rev | TTTAAACAATCTTTCCTGCAACA | qPCR |
| 2336 | cxcl10-fwd | TGGCATTCAAGGAGTACCTC | qPCR |
| 2337 | cxcl10-rev | TTGTAGCAATGATCTCAACACG | qPCR |
| 2245 | shcGAS-fwd | ccggTTAGTTTTAAACAATCTTTCCTctcgagAGGAAAGATTGTTTAAAACTAAtttttg | shRNA |
| 2246 | shcGAS-rev | aattcaaaaaTTAGTTTTAAACAATCTTTCCTctcgagAGGAAAGATTGTTTAAAACTAA | shRNA |
